# Eukaryotic RNA-guided endonucleases evolved from a unique clade of bacterial enzymes

**DOI:** 10.1101/2023.08.09.552727

**Authors:** Peter H. Yoon, Petr Skopintsev, Honglue Shi, Lin-Xing Chen, Benjamin A. Adler, Muntathar Al-Shimary, Rory J. Craig, Zheng Li, Jasmine Amerasekera, Marena Trinidad, Hunter Nisonoff, Kai Chen, Arushi Lahiri, Ron Boger, Steve Jacobsen, Jillian F. Banfield, Jennifer A. Doudna

## Abstract

RNA-guided endonucleases form the crux of diverse biological processes and technologies, including adaptive immunity, transposition, and genome editing. Some of these enzymes are components of insertion sequences (IS) in the IS200/IS605 and IS607 transposon families. Both IS families encode a TnpA transposase and TnpB nuclease, an RNA-guided enzyme ancestral to CRISPR-Cas12. In eukaryotes and their viruses, TnpB homologs occur as two distinct types, Fanzor1 and Fanzor2. We analyzed the evolutionary relationships between prokaryotic TnpBs and eukaryotic Fanzors, revealing that a clade of IS607 TnpBs with unusual active site arrangement found primarily in Cyanobacteriota likely gave rise to both types of Fanzors. The wide-spread nature of Fanzors imply that the properties of this particular group of IS607 TnpBs were particularly suited to adaptation and evolution in eukaryotes and their viruses. Experimental characterization of a prokaryotic IS607 TnpB and virally encoded Fanzor1s uncovered features that may have fostered coevolution between TnpBs/Fanzors and their cognate transposases. Our results provide insight into the evolutionary origins of a ubiquitous family of RNA-guided proteins that shows remarkable conservation across domains of life.

## INTRODUCTION

Insertion sequences (IS) in the IS200/605 and IS607 families occur widely in prokaryotes and encode an RNA-guided endonuclease known as TnpB, the evolutionarily ancestor of type V CRISPR nucleases (Cas12). Analogous to Cas12 enzymes, TnpBs use a guide RNA called right end element RNA (reRNA) or ωRNA to direct cleavage of sequence-complementary DNA molecules (1, 2). However, unlike CRISPR-Cas12, the prevalence of TnpB family proteins extends beyond prokaryotes and their viruses. TnpBs share sequence and structural homology with Fanzor proteins found in eukaryotes including metazoans, fungi, and protists, as well as giant eukaryotic viruses known as nucleocytoplasmic large dsDNA viruses (NCLDV) (3). Like prokaryotic TnpBs, Fanzors occur within transposable elements (TE) that encompass a wide range of transposon families that includes non-prokaryotic IS607 elements (3, 4). Although the biological function of TnpBs and Fanzors is an area of active research, representative proteins within these families have been shown to be RNA-guided DNA endonucleases (1, 2, 4, 5).

Horizontal gene transfer (HGT) plays a crucial role in the evolution of prokaryotes, and mounting evidence indicates that it also exerts a significant influence on the evolution of eukaryotes (6, 7). Notably, despite many examples of HGT from prokaryotes to eukaryotes, prokaryotic IS-like sequences are rare in eukaryotic genomes (8). Nonetheless, IS607 elements represent an unusual instance of horizontally transferred TEs that have become distributed in diverse eukaryotic viruses, and sporadically in eukaryotic genomes (8). As such, IS607 elements provide an unique opportunity to assess features of a prokaryotic IS and their components that facilitate conserved function across the tree of life.

Here we show that an unusual group of IS607 TnpBs with derived nuclease active site signatures likely gave rise to Fanzors. We find Cyanobacteriota as the likely prokaryotic source of Fanzors, and further find evolutionary intermediates between Fanzor1 and Fanzor2, the two types of Fanzor proteins found in eukaryotes. Through functional analysis of the bacterial IS607 TnpB ISXfa1, we report their robust RNA-guided DNA cleavage activity, as well as characteristics that provide insight into the co-evolution of IS607 TnpBs with their transposase. We further find that the guide RNAs for IS607 TnpBs and Fanzors experimentally identified in this study share secondary structural features that underscore their evolutionary relationship. Our findings suggest that beyond the ability to catalyze RNA-guided DNA cutting, some IS607 TnpBs may have properties that made them particularly adaptable to eukaryotic systems.

## MATERIALS AND METHODS

### Metagenomic analysis of TnpB candidates

The ggKbase platform (https://ggkbase.berkeley.edu/) is an analysis tool for metagenomic datasets. All the protein sequences were searched against the Pfam HMM databases of the three TnpB domains, i.e., ‘OrfB_IS605’, ‘HTH_OrfB_IS605’, ‘OrfB_Zn_ribbon’ using hmmsearch (HMMER 3.3 (Nov 2019)) with the parameter of “--cut_nc” (9, 10). The resulting hits with at least one of three domains were identified as TnpB candidates for further analyses. The TnpB candidates with a length range of 350-550aa were clustered based on pairwise protein sequence similarity using MMseqs2, with the parameters as of “--min-seq-id 0.5 --cluster-mode 2 --cov-mode 0 -c 0.5” (11). A total of >140,000 TnpB candidates was identified, which could be further separated into 1,625 different TnpB clusters containing at least five sequences based on their pairwise sequence similarity. The identification of IS605 or IS607 was based on the detection of TnpA next to the TnpB genes, via HMM based search. Once a given TnpA gene was identified with a domain of “Y1_Tnp”, the system was assigned as IS605, if with a domain of “Resolvase”, then as IS607.

### Bioinformatic identification of Fanzor proteins and curation of Fanzor dataset

Non-redundant eukaryotic protein sequences were downloaded from NCBI (November 2022), resulting in 96,197,316 unique sequences. To find putative Fanzor proteins, three HMM models were employed. First, an HMM of the OrfB_Zn_ribbon domain (pfam07282) was used to search the downloaded set of eukaryotic protein sequences using hmmsearch (HMMER 3.3 (Nov 2019)) (9). To filter these hits further we built HMMs on two conserved sub-regions of the RuvC domain (RuvC2 and 3). HMMs of these sub-regions were built by aligning 69 seemingly intact Fanzor sequences from Bao et al., 2013 that were manually selected based on AlphaFold2 structural prediction (12). Two conserved regions were manually identified in the MSA and the resulting HMMs were built using hmmbuild (HMMER 3.3 (Nov 2019)) (9). Sequences that matched the OrfB_Zn_ribbon domain were subsequently searched against the two RuvC sub-region HMMs, again using hmmsearch. Sequences that contained hits to both of these domains were considered putative Fanzors. Next, we extracted the OrfB_Zn_ribbon domain as well as the two RuvC sub-regions from the hits using hmmalign (HMMER 3.3 (Nov 2019)) (9). The extracted regions were used to augment the original alignments of the OrfB_Zn_ribbon and two RuvC sub-regions, where sequences that were hits were added to the original MSA by re-aligning all the sequences using hmmalign and a new profile HMM was built using default hmmbuild parameters. New HMMs were built from these augmented alignments and another round of search was conducted as described above. We continued this process for a total of 5 rounds of searching the eukaryotic protein sequences. Following rounds of manual curation according to ClustalW, where sequences that failed to align at highly conserved residues were filtered, sequence clusters were identified with CLANS and manually enriched with BLASTp (13–15). Finally, the accumulated Fanzor sequences were submitted to AlphaFold2, and the resulting models showing remote structural homology to TnpB were selected as the set of manually curated Fanzors used for phylogenetic and structural analyses.

### Phylogenetic analysis

Before performing multiple-sequence alignments, MMSeqs2 (--min-seq-id 0.5 --cluster-mode 2 -c 0.5 for the large-scale TnpB-Fanzor tree in Figure 1D and Supplementary Figure S1, and --min-seq-id 0.8 -c 0.8 for all other trees in the study) was used to remove highly similar sequences in the dataset (11). The reduced set was then aligned using ClustalW (13). Poorly aligned regions were then filtered with TrimAl (-gt 0.9, -cons 60) (16). Further, IQ-TREE2 was used to generate a maximum likelihood phylogeny with automatic model selection and ultrafast bootstrapping (--bb 1000) (17). The resulting consensus trees were imported to Interactive Tree of Life (iTOL) for post-processing(18).

**Figure 1.**
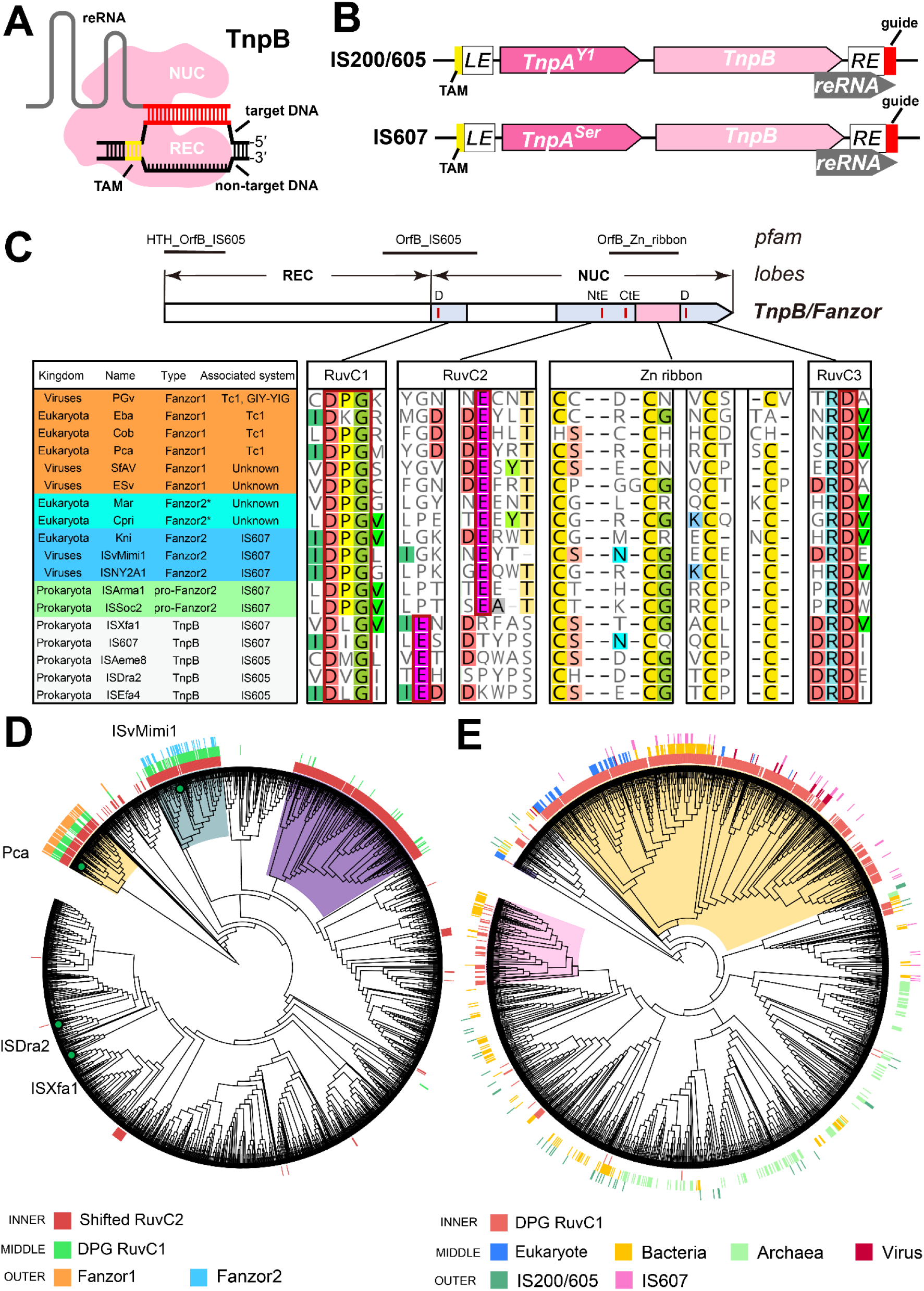
Phylogenetic relationship between TnpBs and Fanzors. A) Cartoon depicting a transposon-associated TnpB nuclease and its guiding reRNA, mediating dsDNA cleavage similar to Type V CRISPR-Cas enzymes. B) A diagram showing typical IS200/605 and IS607 transposons architectures, including the left end (LE) and the right end (RE) of the transposon, the encoded TnpA transposase and the reRNA region at the TnpB. The TnpA-specific domains that differentiate IS200/605 (HUH-transposase: TnpA^Y1^) and IS607 (Serine-recombinase: TnpA^Ser^) are shown. Transposon-associated motif (TAM) and the guide region (similar to spacer for CRISPR-Cas nucleases) are highlighted in yellow and red colors, respectively. C) Sequence signatures of Fanzors and phylogenetically related TnpBs. The right panel depicts MSA of TnpB and Fanzor RuvC subdomains and ZnF motifs. Fanzors and related TnpBs, or pro-Fanzors, sequences share the DPG motif at RuvC1 and the C-terminally shifted catalytic glutamate at RuvC2. The TnpAs and kingdoms associated with the TnpB nucleases are also shown in the table on the left side. Fanzor2* and pro-Fanzor refer to the types of non-IS607 Fanzor2s and Fanzor-like TnpBs that are classified in this study. D) Phylogenetic tree of a representative set of Fanzors and TnpBs fetched from clusters identified in our metagenomics analysis (see Supplementary Figure S1 for the full set). Eukaryotic Fanzor1s are indicated by orange bars in the outer ring, and form one distinct clade (yellow range), whereas Fanzor2 are depicted with blue bars in the second clade (blue-grey range). Both clades share the DPG motif in the RuvC1 and a C-terminally shifted RuvC2 catalytic glutamate, depicted with red and green bars, respectively. Some clades of TnpB may have independently evolved similar features, however, were not associated with eukaryotic counterparts (see e.g. purple range). The tree is re-rooted using Fanzor1 as the outgroup. E) Phylogenetic tree of selected Fanzor2s and TnpBs with C-terminally shifted RuvC2 glutamate residues. Note that DPG appears as a dominant signature motif for Fanzor1s and Fanzor2s, as can be seen in the inner track (red bars). The middle track annotates leaves from NCBI as being either archaeal (light green), bacterial (yellow), eukaryotic (blue), or viral (burgundy). The outer track annotates nodes as having an associated IS200/605 TnpA (dark green) or IS607 TnpA (pink). Yellow range indicates clade containing pro-Fanzor and Fanzor2. Pink range indicates alternative clade showing enrichment of TnpBs sharing RuvC1 and RuvC2 signatures with Fanzors. This tree was re-rooted using Fanzor1 as the outgroup.

### Regular Expression to identify and classify TnpB and Fanzor candidates on NCBI

The Max Planck Institute Bioinformatic Toolkit (MPI) PatternSearch tool (https://toolkit.tuebingen.mpg.de/tools/patsearch) was used to identify TnpB sequences on NCBI NR that shared RuvC1/2 sequence features with Fanzors (19). nr_bac, nr_arc, and nr_euk was searched using D{P}Gx(120,160)ExxxxxxCxxCx(12,18)CxxCx(4,8)D and DPGx(120,160)ExxxxxxCxxCx(12,18)CxxCx(4,8)D as queries. For results from nr_euk, we were able to confirm that this Regular Expression(RegEx) effectively captures the diversity of Fanzor2 sequences we identified through our HMMER and pBLAST searches outlined in the “Bioinformatic identification of Fanzor proteins and curation of Fanzor dataset” section. We further filtered these proteins using NCBI Batch CD tool (https://www.ncbi.nlm.nih.gov/Structure/bwrpsb/bwrpsb.cgi) at default parameters, retaining only hits to guided_TnpB (NF040570), guided_TnpB superfamily (cl45887), HTH_OrfB_IS605 superfamily (cl13722), OrfB_Zn_ribbon (pfam07282), OrfB_Zn_ribbon superfamily (cl37560), InsQ superfamily (cl34004), InsQ (COG0675), IS200_TnpB (NF038281), IS200_TnpB superfamily (cl45733), OrfB_IS605 (pfam01385), OrfB_IS605 superfamily (cl44414). This annotation of motifs was used for phylogenetic trees in Figures 1E and Supplementary Figure S2.

### Annotation of associated transposases

To further annotate the RegEx identified TnpB/Fanzor candidates as being either IS200/605 or IS607-associated, we extracted the amino acid sequences of the immediately neighboring genes based on their NCBI accession ID using a custom Python script. We then examined their PFAMs using NCBI Batch CD tool to determine whether they encode for IS200/605 or IS607 TnpAs (20). Specifically, we filtered for IS200/605 hits using ‘Y1_Tnp’ (pfam01797), ‘Y1_Tnp_superfamily’ (cl00848), and transpos_IS200 (NF033573), and IS607 hits using ‘transpos_IS607’ (NF033518), ‘transpos_IS607 superfamily’ (cl41297), ‘Resolvase’ (cl02788), and ‘Recombinase’ (cl06512). Such annotation was implemented in Figures 1E, and Supplementary Figure S2.

### Identification of prokaryotic Fanzor1 candidates

At first, MPI PatternSearch tool (https://toolkit.tuebingen.mpg.de/tools/patsearch) was used to identify Fanzor1-like sequences using DPGx(120,280)Exxxxxx[CH]xxCx(10,65)Cx(5,50)RDxxxxxN as a query against the nr_bac database, leading to 288 hits (19). As an alternative approach, DPGx(120,280)Exxxxxx[CH]xxCx(10,65)Cx(5,50)RD was used as a query, which yielded 3,192 sequences. Filtering for proteins between 500-1000aa in length retained 20, and 1710 sequences from each dataset, respectively. For the set of 20 sequences, an MSA using curated Fanzor1 sequences was performed to identify proteins that aligned well to the conserved regions not included in the RegEx, which resulted in only one protein sequence. For the set of 1710 sequences, proteins were clustered using MPI MMSeqs2 tool with low stringency (min-seq id 0.1 and coverage .1) to identify proteins that clustered with Fanzor1 proteins (19). This method identified the same protein found in the first more stringent RegEx search. To enrich proteins similar to this mutual hit, a PSI-BLAST search was conducted using NCBI’s webportal (https://blast.ncbi.nlm.nih.gov/Blast.cgi) (21). The search taxa was limited to “bacteria”, and default parameters were used. In each round, only hits larger than 500aa were retained. After three rounds, the search failed to yield any new hits, and was therefore stopped at 16 sequences total, which were further analyzed by manual examination of their contigs. The analysis of selected genes in the contigs with pBLAST revealed homology to proteins found both in bacteria and NCLDV predating on eukaryotes. This ambiguity in the contigs, in addition to the apparent scarcity of intact prokaryotic sequences homologous to Fanzor1, indicated insufficient evidence for HGT of Fanzor1 from bacteria.

### Co-conservation analysis of TnpA and TnpB/Fanzor pairs

For the co-conservation analysis of TnpA and TnpB/Fanzor2 pairs, TnpBs annotated as IS607-associated were parsed from the ISFinder database, and merged with TnpBs with C-terminally shifted glutamate and DΦG motifs, and Fanzor2s, parsed from the ggKbase and NCBI as described above. A total of 197 TnpB/pro-Fanzor/Fanzor2 having TnpA co-encoded in the TE were selected with custom scripts as described above. Such were visualized with CLANS clustering software. TnpBs and Fanzors in four CLANS clusters proximal to ISXfa1 and Fanzor2s were selected for further analysis (a total of 158), together with their associated IS607 TnpA. MSA was performed on sets of TnpB/Fanzors and separately on TnpA with Clustal Omega in Geneious Prime (Geneious 2023.0.4 (https://www.geneious.com)), and the values corresponding to (1) TnpA-TnpA and (2) TnpB-TnpB (Fanzor2-Fanzor2) similarity in the two “% identity” matrices were obtained. These were used to plot the sequence identity histogram of pairs of TnpA and TnpB/Fanzor between two distinct transposable elements (i1 and i2) in MATLAB and further depicted in Supplementary Figure S4.

### Transposon curation

DNA-sequences containing the transposon and flanking sequences (typically 2kb upstream and downstream of the TnpB) were extracted and searched against the assembly using blastn with default parameters. BLAST-hits showing high query coverage (>50%) were extended by 2kb on each end to capture flanking sequences, and then aligned to the query using MAUVE whole genome alignment to identify transposon boundaries and associated features (Geneious 2023.0.1 (https://www.geneious.com)) (15, 22).

### *E. coli* TAM determination assay

The loci of ISXfa1 TnpB overlapping with its reRNA were synthesized (Twist Biosciences) and golden-gate cloned into a vector (Cam resistance) under the control of a single tetracycline-inducible promoter (TetR/pTet) as the Single plasmid (pSingle), used for E.coli TAM depletion assay, E.coli plasmid interference assay and small RNA seq. Alternatively, the ISXfa1 TnpB and its reRNA were individually golden-gate cloned into a vector under two different promoters (TetR/pTet for TnpB and pJ23119 for reRNA) as the Split plasmid (pSplit). The 3′-end of the reRNA were flanked by a HDV ribozyme and the guide region was kept as 20-nt. The target sites with different TAM sequences were golden-gate cloned into the Target plasmid (pTarget) with Amp/Carb resistance.

Electrocompetent *E.coli* DH10β competent cells harboring the TAM library plasmid (Amp/Carb resistance) with six randomized nucleotides (6N) as TAM region which flanks the target site were prepared as follows. The randomized regions were introduced by PCR from the Target plasmid and then golden-gate ligated. The golden-gate products were transformed into electrocompetent *E.coli* DH10β competent cells (NEB) using 0.1 mm electroporation cuvettes (Bio-Rad) on a Micropulser electroporator (Bio-Rad). The cells were plated on LB-Agar plate with 100 μg/mL of Amp. Over 100,000 colonies were scraped to ensure appropriate coverage of all possible combinations of the randomized TAM region. The cells were then inoculated 1:100 into a 1 L LB culture with 100 μg/mL of Amp at 37 °C to OD_600_=0.6. The cells were then cooled to 4 °C, pellet and sufficiently washed by ice-cold water and 10% glycerol. The pellets were then resuspended in 2 mL 10% glycerol, aliquoted into 50 μL and stored in -80 °C.

These TAM library competent cells were then thawed and transformed with 100 ng of the pSingle (Cam resistance) using 0.1 mm electroporation cuvettes (Bio-Rad) on a Micropulser electroporator (Bio-Rad). The cells were recovered in 1 mL LB culture at 37 °C for one hour. After recovery, the culture was inoculated 1:100 into 3 mL LB culture with 34 μg/mL of Cam, 100 μg/mL of Carb (equivalent to Amp) and was induced with 200 nM aTc at 26°C for 24 hours. The plasmids that had been propagated were then isolated using the QIAgen Miniprep Kit (Qiagen). TAM motifs that were preferentially depleted were identified using amplicon sequencing of the TAM library plasmid. The library was sequenced on an Illumina NovaSeq by The High-Throughput Sequencing Core in UCLA Broad Stem Cell Research Center, with a 150x150 paired-end run configuration and average depth of 500,000 reads per sample. TAM preferences were characterized from FASTQ files using a custom Python script. Briefly, six-nucleotide TAMs were extracted and counted. TAM frequencies were then normalized to the sequencing depth of each sample. Sequence logos were generated with Logomaker (version 0.8) using TAMs with a log2 fold-change of 5 or greater relative to the original plasmid library (23).

### *E. coli* plasmid interference assays

Electrocompetent *E.coli* DH10β competent cells were double-transformed with 100 ng of the pSingle or the pSplit (Cam resistance) and 100 ng of the pTarget (Amp/Carb resistance) using 0.1 mm electroporation cuvettes (Bio-Rad) on a Micropulser electroporator (Bio-Rad). The cells were recovered in 1 mL LB culture at 37 °C for one hour. After recovery, 5-fold dilution series of the recovery culture were prepared and 5 μL of the dilution were spotplated on double-antibiotic plates (LB-Agar plate with 34 μg/mL of Cam, 100 μg/mL of Carb and 2 nM aTc) as well as single-antibiotic control plates (LB-Agar plate with 34 μg/mL of Cam and 2 nM aTc). For plates with no visible colonies, 400 μL of the 1 mL recovery culture were all plated on the double antibiotic plate for improving detection limit. The plasmid interference efficiency was quantified by the normalized colony-forming units (Norm. CFU) which represents the CFU on the double-antibiotic plate (Carb+Cam) divided by the CFU on the single-antibiotic control plate (Cam).

### *E. coli* RNA expression and extraction

The pSingle containing ISXfa1 TnpB and reRNA was transformed into E.coli DH5α competent cells (Thermo Fisher Scientific). A single colony was picked and inoculated in 3 mL LB culture with 34 μg/mL of Cam and 100 μg/mL of Carb in 37 °C overnight as starting culture. The starting culture was then inoculated 1:100 in 3 mL LB culture with 34 μg/mL of Cam, 100 μg/mL of Carb and 200 nM aTc to induce the RNA expression. Once the OD_600_ reaches 1.0, 400 μL of the cells were taken and mixed with 800 μL of RNAprotect Bacteria Reagent (QIAgen). After 15 min, the cells were pellet by centrifuge, and stored in -80°C after removing all the supernatant. The next day, the pellets were resuspended in 98% formamide and 10 mM EDTA and lysed at 70 °C for 10 min. The total RNA were then extracted from the lysate using Zymo Quick-RNA Miniprep Kit (Zymo Research) and used for small-RNA sequencing library preparation.

### Insect cell experiments

Expi-Sf9 (GIBCO) cells were cultured in Expi-CD medium (GIBCO) in Polycarbonate Erlenmeyer Flasks with a vent cap (Corning). Cultures were kept in a shaking incubator (New Brunswick) at 26 °C and 100 RPM with a 50mm shaking diameter. Cells were maintained at densities between 5x10^5^ and 5x10^6^, with the volume of the liquid culture remaining below 1/5th of the total volume of the flask. Cell viability and density were measured using Trypan-Blue (GIBCO) on a CytoSMART Cell Counter (Corning).

Bacmids were generated by cloning in DNA fragments into a pFASTBAC vector through PCR amplification of the vector and insertion of synthesized gene-fragments (Twist Biosciences) via Golden Gate Assembly, using DH5a cells (ZYMO), and transforming the cloned plasmids into MAX EFFICIENCY DH10BAC (GIBCO) or EmBacY (Geneva Biotech) *E. coli* strains following manufacture protocols. Chemically transformed cells were allowed to recover for 4 hours in 2XYT media, and plated on a Luria-Bertani (LB)-Agar plate containing kanamycin (50 μg/ml), gentamicin (7 μg/ml), tetracycline (10 μg/ml), IPTG (40 μg/ml) and X-gal (100 μg/ml). After 48 hours at 37 C, large white colonies were picked and cultured in liquid culture containing kanamycin (50 μg/ml), gentamicin (7 μg/ml), tetracycline (10 μg/ml). Bacmids were purified by lysing *E. coli* using the ZymoPURE Miniprep kit (ZYMO), followed by ethanol precipitation for isolation of bacmid DNA. Transgene insertion into the bacmid was confirmed via PCR using M13 reverse primer and a primer binding to the transgene.

12.5 μg of freshly prepared bacmid was transfected into a 25mL culture containing 62.5x10^6^ Expi-Sf9 cells using ExpiFectamine™ Sf Transfection Reagent following manufacturer instructions (GIBCO). Supernatant containing P0 baculovirus was harvested 3 days post-infection by using centrifugation to separate from cells, and titered using the baculovirus titering kit (Expression Systems) and Attune NxT Flow Cytometer with an autosampler. P0 virus was used directly to infect a 25 mL culture with 5x10^6^ Expi-SF9 cells at an MOI of 5. Cells were harvested 3 days post-infection for downstream analysis.

### Small RNA sequencing

Total RNA was extracted using a one-step hot formamide extraction method for all samples, where cells are incubated in a 18 mM EDTA and 95% formamide solution at 65 °C for 5 minutes (24). Cell debris was removed by centrifugation, and 2 volumes of RNA binding buffer (ZYMO) was added to each volume of RNA sample. 1 volume of ethanol was then added to the mixture, and loaded to a Zymo-Spin IIC column. Following on-column DNAse treatment, total RNA was eluted using water. ∼200 ng of total RNA was used for rRNA depletion reactions using NEBNext rRNA kit (bacteria) following manufacturer protocols. For eukaryotic samples, custom probes designed using NEBNext Custom rRNA depletion tool were used. The rRNA depleted RNA was purified using a Zymo-Spin IIC column, treated with T4 PNK using T4 PNK buffer (NEB) at 37 C for 30 minutes before spiking in ATP and further incubating. Further end treatment was performed using mRNA Decapping Enzyme (NEB) pre-mixed with its buffer directly into the PNK reaction and incubating at 37 C for 30 minutes. End-repaired RNA was purified using a Zymo-Spin IIC column, and used for Collibri small-RNA-seq library preparation reactions following manufacturer protocols (INVITROGEN). The resulting NGS adapter ligated cDNA was then amplified with KAPA HiFi HotStart ReadyMix using NGS indexing primers for 25 cycles (ROCHE). Amplified DNA fragments between 150∼500nt were size selected by gel extraction purification following gel-electrophoresis in a 4% SYBR-GOLD E-GEL. Before loading into the sequencer, library concentrations were quantified using KAPA-QUANT qPCR kit (ROCHE). NGS reads were merged using BBMerge using a normal merge rate and then mapped onto the TnpB containing locus using Geneious mapper at Low Sensitivity to identify reRNA boundaries (Geneious 2023.0.1 (https://www.geneious.com)).

### Rearing and virus infection of *S. frugiperda* larvae

Insect hemolymph containing budded *Spodoptera frugiperda* ascovirus 1a (SfAV) vesicles was received as a generous gift from Brian Federici (University of California, Merced). *Spodoptera frugiperda* larvae were grown on an artificial diet (Benzon Research) at room-temperature until third-instar stage, when they were pricked with a needle contaminated with SfAV containing hemolymph. 5 days post treatment, infected larvae with opaque hemolymph were selected for RNA isolation. Both isolated hemolymph and whole larvae homogenized using liquid nitrogen and mortar and pestle were used for RNA-seq experiments. For *in vivo* protein expression experiments using recombinant baculovirus, larvae were infected by injecting with baculovirus containing clarified culture media and homogenized using the same technique as above.

### Western blots

Homogenized larval samples were prepared in buffer containing 50 mM Tris-HCl, 500 mM NaCl, and 5 mM TCEP on ice. Homogenized larval samples were further sonicated in 1.5ml Eppendorf tubes at 60% power setting, 30 s on/30 s off for a total sonication time of 5 min at 4 °C. Large particulates from this solution were filtered using a funnel layered with Kimwipes prior to centrifugation. After removing debris by centrifugation at 16,000 g for 10 min, protein concentration in the supernatant was measured on a nanodrop spectrophotometer. ∼50 mg protein lysate was used as stock that was diluted 5-fold per serial dilution in each well. Lysate and dilutions were denatured in 1x Laemmli buffer at 95C for 30 seconds and resolved by SDS-PAGE on a 8-20% gradient gel (BioRad). Following trimming of the gel, proteins were electro transferred to a PVDF membrane activated in methanol for 1 min. The membrane was blocked with blocking buffer (PBS/0.05% Tween-20 containing 5% milk) for 1 hr, incubated with primary antibody in blocking buffer for 1 hr, washed three times with PBS/0.05% Tween-20 for 5 minutes each, incubated with secondary antibody conjugate in blocking buffer for 1 hr, and washed three times again with PBS/0.05% Tween-20 for 5 min each. Protein bands were visualized using LI-COR Odyssey CLx with Image Studio v5.2 software at 700/800 nm channels.

## RESULTS

### An unusual class of RNA-guided TnpB endonucleases generated eukaryotic Fanzors

The IS200/605 and IS607 families of transposons encode proteins TnpA and TnpB, and a non-coding RNA called reRNA. TnpB is an RNA-guided endonuclease that uses reRNA for DNA target recognition and interference (Figure 1A) (1, 2). Although both IS200/605 and IS607 elements share the TnpB gene, their TnpA transposases are unrelated (Figure 1B). Whereas IS200/605 elements encode a Y1-transposase (TnpA^Y1^) that mediates “peel-and-paste” transposition of single stranded DNA, IS607 elements encode a serine-recombinase (TnpA^Ser^) that inserts double-stranded DNA into short dinucleotide motifs (25–29). Whereas IS607 elements are found in all three domains of life, IS200/605 elements have not yet been reported to occur in eukaryotes despite their abundance relative to IS607 elements (30).

Fanzors, the eukaryotic homologs of TnpBs, are categorized into two types: Fanzor2 that closely resembles prokaryotic TnpBs and found in IS607 elements, and Fanzor1 that is divergent and found in various eukaryotic transposon families (3). To determine the evolutionary origins of Fanzors we closely evaluated their protein sequence features. TnpBs and their relatives, including Fanzors, share an evolutionarily conserved RuvC nuclease domain whose DED catalytic triad is split into three discontinuous parts, or subdomains, in the primary sequence (RuvC1-3) (Figure 1C). Additionally, many TnpBs and Fanzors contain a C-terminal Zinc-Finger (ZF) motif, also referred to as Zinc ribbon (ZR) domain, that is flanked by the RuvC2 and RuvC3 subdomains (Figure 1C). Multiple sequence alignment (MSA) showed that both Fanzor1s and Fanzor2s contain a highly conserved DPG motif in the RuvC1 subdomain (Figure 1C). Fanzors also contain the previously reported C-terminally shifted RuvC2 glutamate residue, which we find is only six residues away from the ZF motif (Figure 1C) (4). These features are distinct from previously reported TnpB sequences that consist of a RuvC1 region that typically contains a DΦG motif (Φ is the hydrophobic residues I, L, F, W, Y and M), and a RuvC2 region whose active site residue is ∼50 residues upstream of the ZF motif (Figure 1C).

To identify TnpBs that share these features with Fanzors, we first identified >140,000 TnpB candidates from ggKbase metagenomic sequences (see Methods). A Regular Expression (RegEx) that filters for TnpBs with a DPG RuvC1 motif and a C-terminally shifted RuvC2 glutamate residue yielded 1,227 sequences (Supplementary Table S1). We also used RegEx to search for TnpB sequences that showed only one of the two features. This revealed that 5,053 sequences contain only a C-terminally shifted RuvC2 glutamate residue, and that 171 sequences contain only a DPG RuvC1 motif (Supplementary Table S1). In summary, our findings suggest that the combination of the sequence features we identified in both classes of Fanzors are rare, being present in less than 1% of TnpBs in our dataset.

We next explored the evolutionary relationship between TnpBs and Fanzors via phylogenetic analysis. From the set of >140,000 TnpB candidates, we identified 1,625 different TnpB clusters containing at least five sequences based on their pairwise sequence similarity (see Methods for details). We pooled representatives from each cluster and generated a maximum-likelihood phylogenetic tree (see Methods for details; the full phylogenetic tree in Supplementary Figure S1). We found that Fanzor1s formed a single clade (Figure 1D, yellow range). Similarly, Fanzor2s and TnpBs with Fanzor-like sequence features (that is, both the RuvC1 and RuvC2 signatures) formed a well supported second clade (Figure 1D, blue-grey range). Our analysis also identified a large clade of TnpBs that may have evolved a shifted RuvC2 glutamate residue independently to Fanzors (Figure 1D, purple range). This clade of TnpBs, however, is differentiated by the fact that it typically lacks the RuvC1 DPG motif found in Fanzors (Figure 1D, middle green track).

To gain greater resolution into the evolutionary origins of Fanzors, we performed phylogenetic analysis using a filtered set of TnpB sequences that contain the shifted RuvC2 feature. In this analysis, we also included Fanzor/TnpB sequences from NCBI to enable taxonomic interpretation (see Methods for details), as the short length of most TnpB containing metagenomic contigs made high-confidence taxonomic assignment challenging. The full phylogenetic tree can be found in Supplementary Figure S2. In line with our previous analysis, we found that the vast majority of TnpBs that share both RuvC1 and RuvC2 features with Fanzors were closely related to each other, forming a well supported clade with all Fanzors used in this analysis (Figure 1E, yellow range). Beyond the Fanzor-containing clade, TnpBs with Fanzor-like RuvC1 signatures were enriched to a lesser extent in one additional clade (pink range), but otherwise found sporadically throughout the tree (Figure 1E). Broadly, we found that prokaryotic IS607 TnpBs in this clade were found almost exclusively in Cyanobacteriota (Figure 1E, yellow middle track within the yellow range; also see Supplementary Table S2). Corroborating the relationship between this clade of Cyanoberiota-derived IS607 TnpBs and Fanzor2s, we also found their TnpAs to be related (Supplementary Figures S3, S4). Collectively, our analysis suggests that Fanzor2 emerged from HGT of an unusual type of IS607 TnpBs with divergent active site signatures found primarily in Cyanobacteriota. From here on, we will refer to this type of TnpB as pro-Fanzors, as they represent the putative prokaryotic ancestors of Fanzors.

### Fanzor1 likely evolved from Fanzor2 in eukaryotes

To better understand the evolutionary relationship between the two types of Fanzors, we performed phylogenetic analysis using only the RuvC containing region of the proteins. This analysis clearly separated Fanzor1 and Fanzor2 as two separate clades (Figure 2A and Supplementary Figure S5). The apparently distant phylogenetic relationship between Fanzor1 and Fanzor2 led us to investigate the possibility that the two types of Fanzors evolved from separate HGT events of different types of TnpBs. To do so, we searched for Fanzor1 homologs in the prokaryotic NR database (see Methods section). This search generated a list of 16 candidates, which showed good overlap with the list of bacterial proteins previously reported to be closely related to Fanzor1 (5). Among these 16 proteins, all of which were identified by metagenomics, only 4 were encoded on contigs longer than 5kb. BLASTp queries of the genes neighboring the Fanzor1 candidates revealed that in all four instances, the Fanzor flanking genes were a mix of bacterial and nonbacterial genes belonging to eukaryotic giant viruses and eukaryotes. This suggests that these candidates are either genome assembly artifacts, or proteins recently acquired via HGT from a eukaryote or their viruses. In both cases, it would appear that established Fanzor1-like proteins are extremely rare to non-existent in prokaryotes.

**Figure 2.**
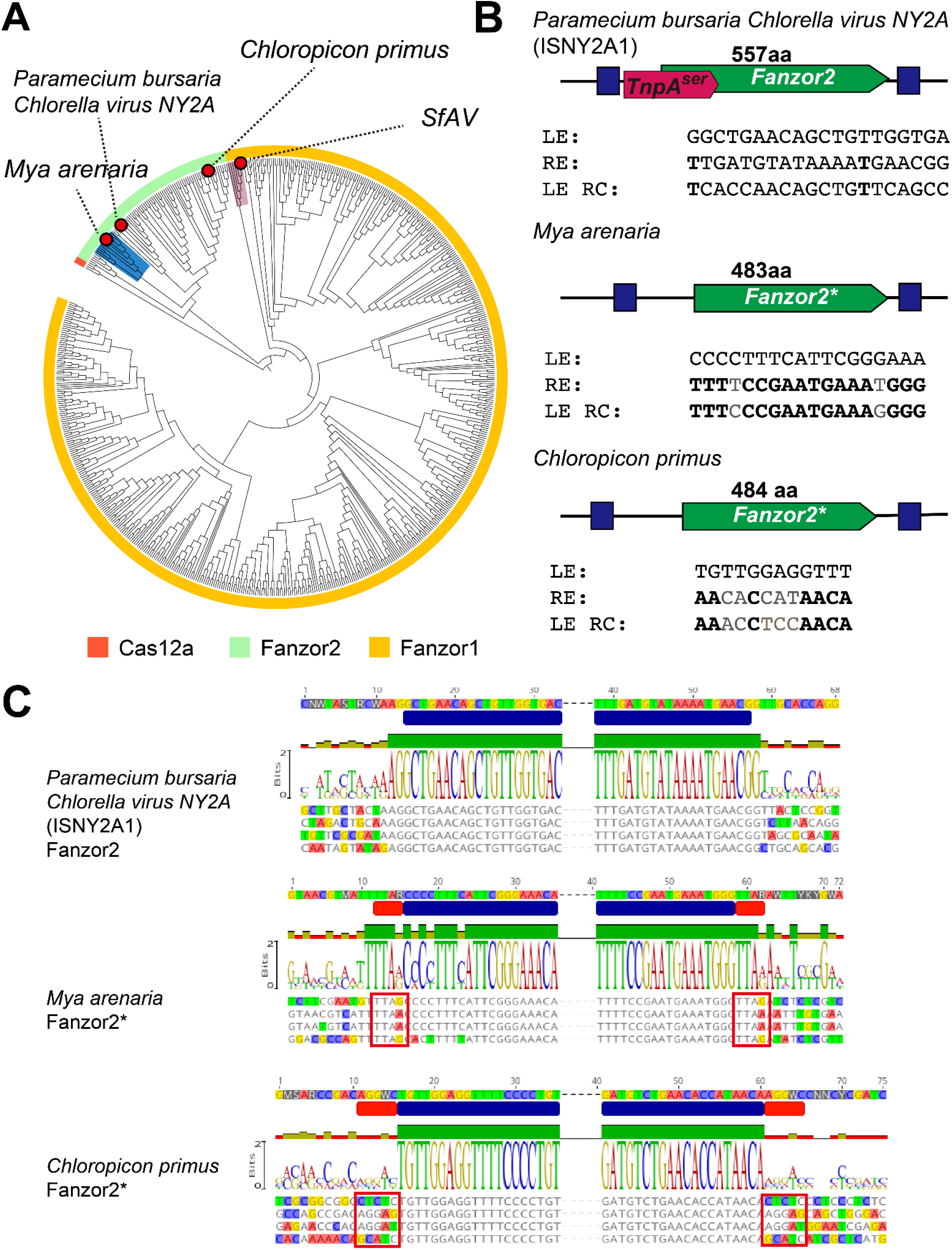
Fanzor2s have been captured by non-IS607 transposons on multiple occasions. A) Phylogenetic tree of Fanzor RuvC sequences. Blue range indicates clade of Fanzor2s found in molluscs and their viruses that has been co-opted by a non-IS607 transposon. Pink range indicates clade of Fanzor1s found in lepidopterans and their viruses, including S. frugiperda Ascovirus (SfAV). Track annotates nodes as being either a Cas12a (red) Fanzor2 (green) or Fanzor1 (yellow). Tree was re-rooted using Cas12a as an outgroup. B) Locus organization of ISNY2A1 Fanzor2 and *Mya arenaria*, and *Chloropicon primus* Fanzor2* encoding elements and their LE/RE (blue boxes). In sequence labels, LE is left end, RE is right end, and LE RC is left end reverse complement. ISNY2A1 locus encodes an IS607 TnpA and Fanzor2, whereas *Mya arenaria* and *Chloropicon primus* loci only encode the Fanzor2*. C) MSA of ISNY2A1, *Mya arenaria*, and *Chloropicon primus* Fanzor2 encoding elements demonstrating transposon boundaries (blue annotation) as well as clear TSD (red annotation and red box) in *Mya arenaria* and *Chloropicon primus*. *Mya arenaria* and other mollusc Fanzor encoding elements have 4nt TSD conserved as TTAN, whereas *Chloropicon primus* has 5nt TSD that shows high degree of variability.

A defining feature of Fanzor1 is that it is found in a wide range of canonical eukaryotic transposon families. This is in contrast to Fanzor2, which previously has been suggested to be only encoded in eukaryotic IS607 elements (3). However, in our analysis of Fanzor2 encoding transposons, we found evidence of Fanzor2s from the alga *Chloropicon primus* and various mollusc species including *Mya arenaria* having been captured by non-IS607 transposons (Figure 2B and C). In contrast to canonical Fanzor2s and their closely related IS607 TnpBs, these Fanzors are encoded by non-autonomous transposons with terminal inverted repeats (TIR) that cause target site duplications (TSD) upon insertion (Figure 2B and C). These features would not be expected of IS607 elements such as ISNY2A1, an IS607 element encoded in the genome of *Paramecium bursaria chlorella virus*, a eukaryotic virus infecting algae (Figure 2B and C). The presence of both TIRs and TSDs suggest that these eukaryotic elements are canonical *cut-and-paste* DNA transposons, although we were unable to identify autonomous partners in the respective genomes to classify them to specific superfamilies. Interestingly, phylogenetic analysis suggested that the algal Fanzor2 is distantly related to the mollusc Fanzor2s, suggesting independent capture of IS607 Fanzor2s by eukaryotic transposons (Figure 2A). The scarcity of Fanzor1 homologs in prokaryotes, combined with the observation of Fanzor2s encoded in eukaryotic DNA transposons suggest an evolutionary scenario where Fanzor1s evolved from ancestral Fanzor2s, which involved Fanzor2 capture by different eukaryotic transposons. From here on, we will refer Fanzor2s encoded in eukaryotic DNA transposons as Fanzor2* to differentiate them from canonical IS607 encoded Fanzor2.

### IS607 TnpB shows features that co-evolved with its serine-recombinase TnpA

To gain insight into the aspects of IS607 elements that may have contributed to their HGT, we next sought to experimentally characterize an IS607 TnpB. We chose to study a TnpB, which shows typical active site arrangement, encoded in an IS607 element found in the *Xylella fastidiosa* genome (hereafter, ISXfa1) (Figure 3A). Small-RNA sequencing of transcripts isolated from *E. coli* overexpression of ISXfa1 TnpB through the right end showed that as reported for both TnpBs, ISXfa1 TnpB associates with a ∼150 nucleotide reRNA that extends ∼17 nucleotides beyond the transposon boundary (Figure 3B) (31).

**Figure 3.**
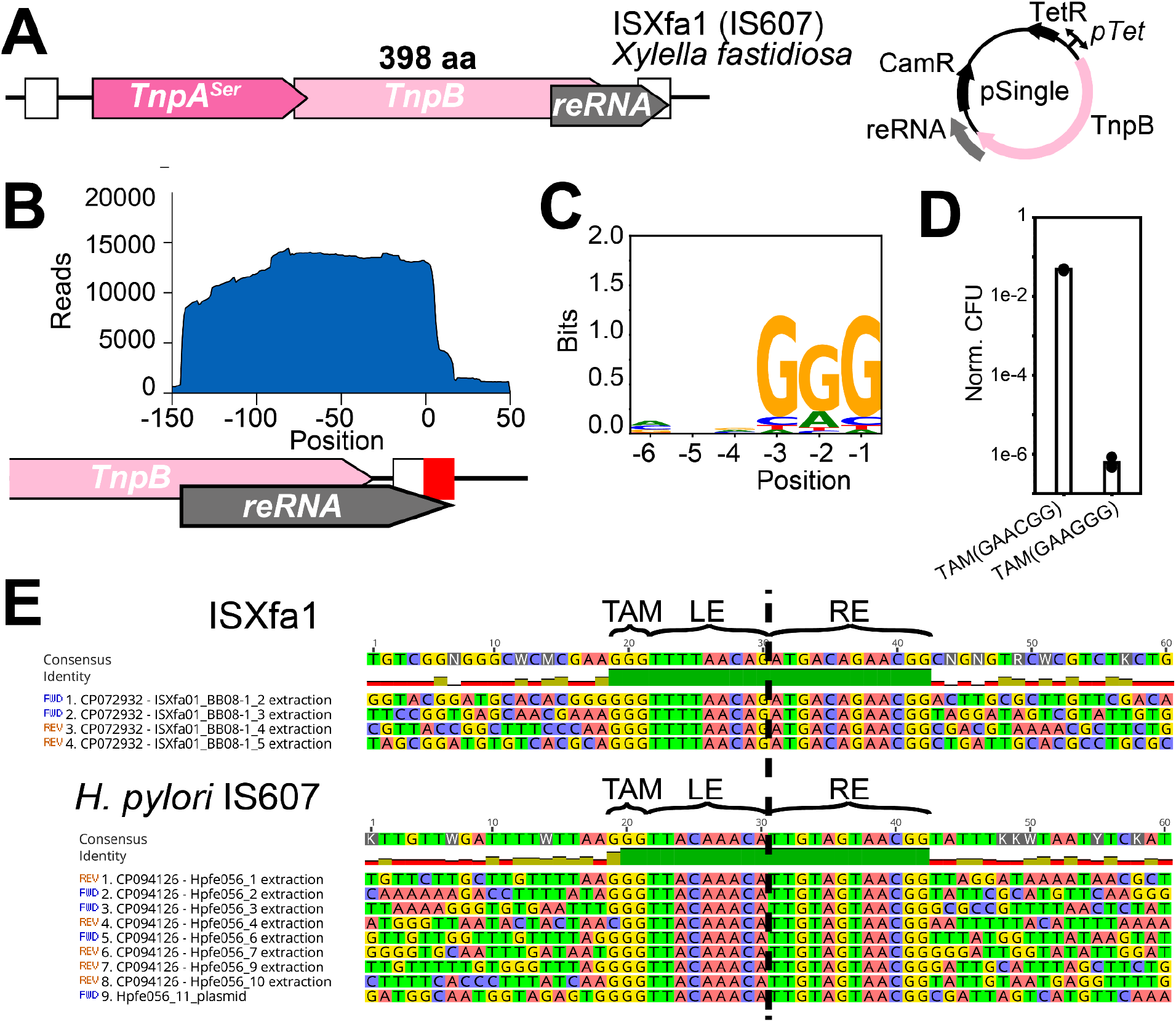
IS607 TnpB is an RNA-guided DNA endonuclease. A) (Left) Cartoon showing domain compositions and reRNA coding sequence within ISXfa1 transposon. (Right) pSingle (Cam resistance) encoding both ISXfa1 TnpB and its reRNA under the same promoter (pTet/TetR). B) Small RNA seq results suggest that the reRNA of ISXfa1 is composed of 144-nt scaffold and 17-nt guide. C) TAM assay showing the preferred TAM sequence of ISXfa1 TnpB. D) Quantification of E. coli plasmid interference assay showing that ISXfa1 TnpB effectively targets a GGG TAM, but not a CGG TAM. E) MSA of IS607 elements in *Xylella fastidiosa* (ISXfa1) and *Helicobacter pylori* (IS607) genomes and plasmids demonstrating conservation of GGG sequences at the insertion site.

We next evaluated whether ISXfa1 TnpB can mediate plasmid restriction given a matching guide and transposon associated motif (TAM). To test this, we utilized a plasmid library depletion assay, which revealed a depletion of GGG motifs when a matching guide was used (Figure 3C). As IS607 transposons typically insert via recombination between matching dinucleotide motifs, the first nucleotide of the transposon right-flank is fixed (27). To determine whether this region was reprogrammable as in the case of IS200/605 TnpBs, we deleted bases in this region and replicated the plasmid depletion assays. Upon deletion of a single G, we observed a depletion of GG motifs, whereas deletion of two G’s lead to a loss of function (Supplementary Figure S6). These results suggest that IS607 TnpBs maintain the same degree of reprogrammability as IS200/605 TnpBs by incorporating the first nucleotide of the right flanking sequence as part of the RNA scaffold.

Previous studies of the IS607 TnpA from *Helicobacter pylori,* which shares 80% sequence identity with ISXfa1 TnpA, inserts into a GGG or CGG sequence when heterologously expressed in *E. coli* (28, 29). In IS200/605 systems, the insertion specificity of the TnpA perfectly matches the TAM of the TnpB (1, 30, 32). We therefore assayed ISXfa1 TnpB for its ability to cleave DNA targets bearing either a GGG or CGG motif using a bacterial plasmid interference assay (Supplementary Figure S7). We found that ISXfa1 TnpB mediates robust interference of plasmids containing a GGG TAM, but not a CGG TAM (Figure 3D, Supplementary Figure S7 and S8). This is in line with the observation that the ISXfa1 reRNA guide region is flanked by a CGG sequence, as its recognition would lead to interference of intact transposons (Figure 3E). We further observed that most IS607 copies in both the *Xylella fastidiosa* and *Helicobacter pylori* genome have been inserted into GGG sequences (Figure 3E). These observations suggest that in IS607 elements, TnpBs may influence the insertion profiles of the transposon by selecting for TAM bearing insertion sites following promiscuous insertion by the TnpA.

### Conservation of guide RNA secondary structure among TnpBs and Fanzors

Cryo-EM structures of the ISDra2 TnpB ternary complex revealed that the core of the reRNA is formed through tertiary interactions including pseudoknot (PK) interactions between a few nucleotides flanking the guide and the apical loop of the stem-loop 3 (Figure 4A) (33, 34). Similar PK interactions have also been observed in Cas12f ternary structures (35, 36). RNA folding analysis of ISXfa1 reRNA suggests that its overall architecture may be conserved with ISDra2 reRNA, with three predicted stem-loops and a potential PK forming region (Figure 4A). RNA-folding analysis of putative reRNA encoding regions of other IS607 TnpBs, including Fanzors and their closely related prokaryotic ancestors, suggested that these features may be generally conserved among IS607 TnpB and Fanzor2s (Figure 4A).

**Figure 4.**
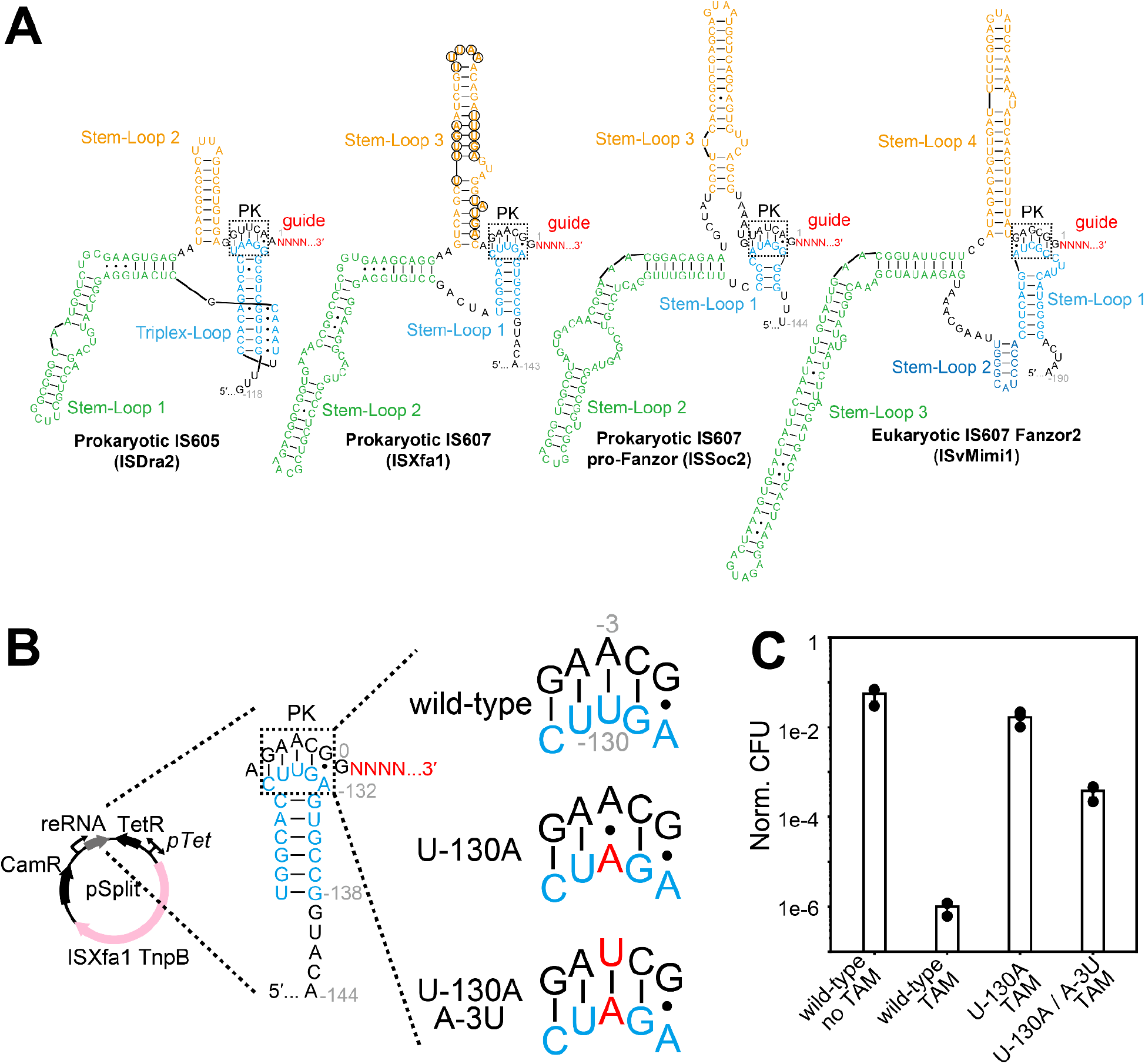
Conservation of reRNA secondary structure between IS605/607 TnpB and Fanzor2. A) Putative secondary structure comparison of reRNA between IS605, IS607, pro-Fanzor IS607, Fanzor2 reRNA. Putative pseudoknot (PK) interactions are highlighted in black dashed box. Regions in the Stem-Loop 3 in the ISXfa1 reRNA corresponding to the TnpA direct repeats are circled in black. B) Compensatory experiments testing the PK interactions in ISXfa1 reRNA. In the pSplit, the TnpB and its reRNA are under different promoters. Two reRNA mutants were designed: The U-130A mutant results in an A-A mismatch in the PK region whereas the U-130A/A-3U with an additional mutation restores the A-U Watson-Crick base pair in the PK. C) The PK interactions are confirmed by E.coli plasmid interference assay as the U-130A mutant disrupted the ISXfa1 TnpB activities while the U-130A/A-3U mutant rescue the TnpB activity

To experimentally validate the reRNA structure, we first destabilized the PK interactions by point mutating U-130 to A-130 at the apical loop in Stem-loop 3 that results in an A-A mismatch instead of an A-U Watson-Crick base pair in the PK (Figure 4B). Compared to the wild-type, the PK destabilizing mutant showed substantially reduced activity (< 10^-4^) in an *E.coli* interference assay (Figure 4C). We then introduced a second rescue point mutation at the guide flanking region (A-3U) to restore a U-A Watson-Crick base pair in the PK. The PK rescue mutants restored the activity by ∼100 fold (Figure 4C). The incomplete rescue could be attributed to the potential contacts between the apical loop in Stem-loop 3 with ISXfa1 TnpB, as indicated in the ISDra2 TnpB cryo-EM structures (33, 34). These results demonstrate that a PK evolutionarily conserved between TnpBs and Fanzor2s is required for the activity of ISXfa1 TnpB.

### Fanzor1 protein and reRNA expression in a native insect host

We next sought to understand how TnpB-family proteins adapt to non-IS607 transposons, as observed in Fanzor2*s or Fanzor1s. We chose to study a Fanzor1 from *Spodoptera frugiperda* Ascovirus 1a (SfAV), a large DNA virus, or NCLDV, infecting lepidopterans that is transmitted via parasitoid wasps (3, 37, 38). The Fanzor-encoding element of SfAV is a simple non-autonomous transposon present at two copies in the genome that only encodes a 606-residue Fanzor1 protein (Figure 5A). This group of Fanzors appears to be encoded in an unknown DNA transposon with short TIRs that inserts into a TTTAN sequence, causing a TTAN target site duplication upon insertion (3).

**Figure 5.**
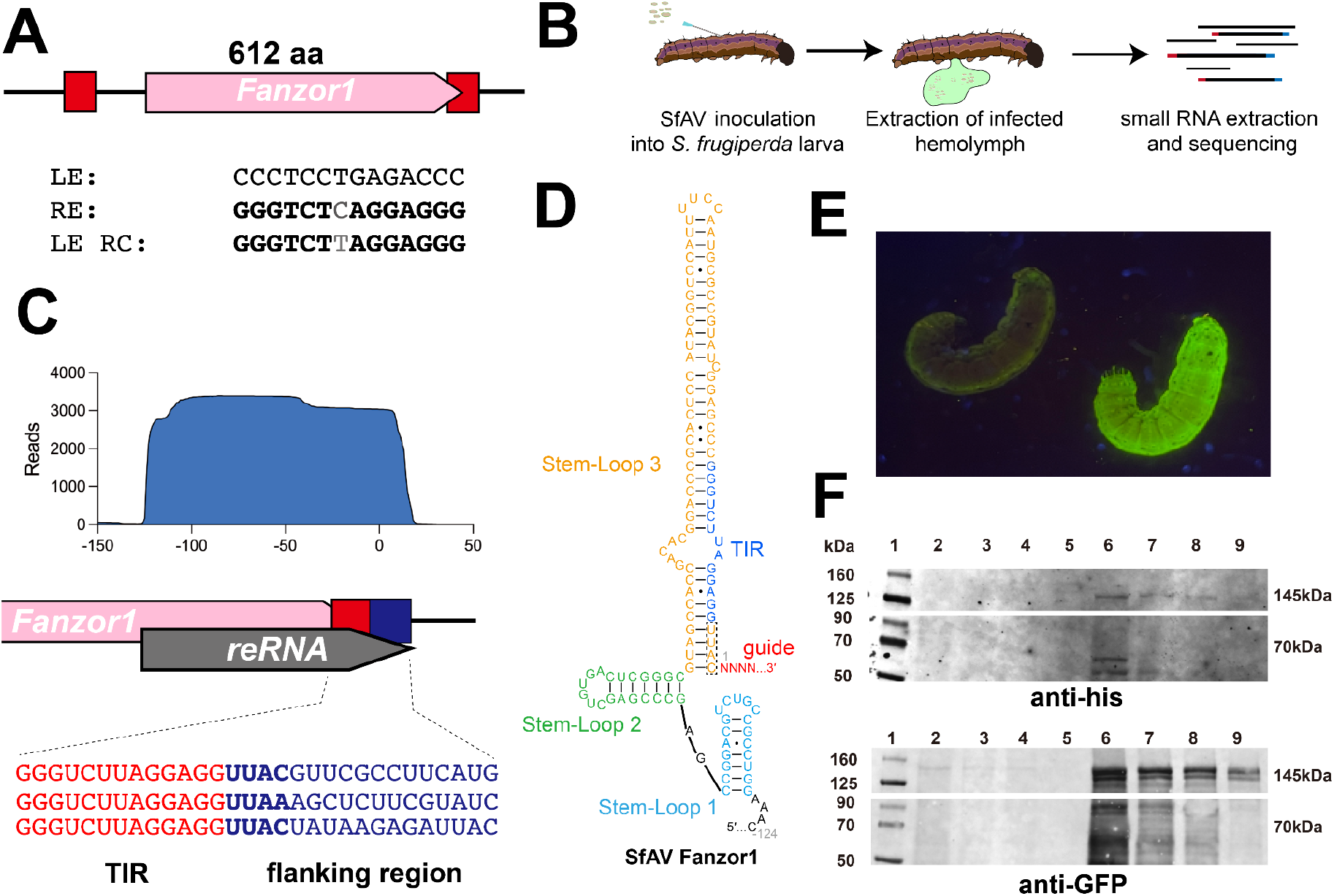
Fanzor1 reRNA and protein expression in native insect hosts. A) Locus organization of SfAV Fanzor1 and its TIR. LE is left end, RE is right end, and LE RC is left end reverse complement. B) Cartoon depicting workflow of *in vivo* SfAV experiments in *S. frugiperda* larvae used to assay reRNA expression. C) Mapping of small RNA-seq reads onto the reference locus encoding SfAV Fanzor1. Reads suggest that the reRNA of SfAV Fanzor1 is composed of ∼120-nt scaffold and ∼20-nt putative guide region that is variable depending on the transcript. D) Putative secondary structure of SfAV Fanzor1 reRNA. Each stem-loop is color coded as depicted in the panel, except for the dark blue nucleotides, which corresponds to the TIR of the Fanzor encoding transposon. E) Picture of *S. frugiperda* larvae infected with recombinant baculovirus encoding his-tagged Fanzor (left) and his-MBP and GFP-CL7 tag fused Fanzor (right). F) Western blot showing expression of SfAV protein in vivo using anti-his and anti-GFP primary antibodies. Lane 1: Chameleon ladder. Lanes 2-5: his-SfAV lysate dilutions. Lanes 6-9: his-MBP-SfAV-CL7-GFP lysate dilutions.

We performed small RNA-sequencing experiments to examine reRNA expression *in vivo* by infecting *S. frugiperda* larvae with SfAV. To do so, we pricked the *S. frugiperda* larvae with an SfAV contaminated needle to mimic oviposition of parasitoids, which effectively transmits the virus (Figure 5B). As ascoviruses primarily replicate in the hemolymph, we isolated and sequenced small-RNAs from hemolymph samples from infected *S. frugiperda* larvae (Figure 5B) (37). Mapping of small-RNA reads to the SfAV genome revealed that the reRNA of the Fanzor was one of the most highly transcribed small RNAs in the SfAV genome (accession: AM398843) (Supplementary Figure 9A). Consistent with previously reported kinetics of ascovirus infection, we found that reRNA abundance increased 5 days post infection (dpi) compared to 2 dpi, when infected larvae began to develop a cloudy hemolymph caused by budded viral vesicles (Supplementary Figure S9A) (39). In both samples, we observed expression of an RNA that mapped to the right end (RE) of the transposons and extended further downstream into the flanking DNA sequence (Figure 5C). Interestingly, we found three different putative guide sequences in our RNA-sequencing data, suggesting the presence of a third unreported copy of the SfAV Fanzor in the genome (Figure 5C).

In addition to assaying expression *in vivo*, we generated recombinant baculovirus encoding SfAV Fanzor and two additional insect virus encoded Fanzor1s and infected Expi-Sf9 cells. Similar to *in vivo* expression, we observed the expression of an reRNA that extended ∼20nt beyond the RE of the transposon in all three samples (Supplementary Figure S9B). RNA-folding analysis of insect viral Fanzor1 reRNA showed that these Fanzor1 reRNAs fold into a 3-stem shape whose global architecture was similar to TnpB and Fanzor2 reRNAs (Figure 4A, 5C and Supplementary Figure S9C). Notably, we found the conservation of a stem-loop flanking the putative guide region in all instances (Figure 4A). In the insect Fanzors, the TIRs and the TSDs formed a part of the conserved stem-loop, suggesting that both are incorporated into the scaffold. The inclusion of the TSD, whose sequence is fixed, into the Fanzor1 reRNA scaffold parallels the inclusion of the transposon flanking nucleotide, whose sequence is also fixed, into the scaffold in ISXfa1 reRNA (Figure 5F). This suggests that TnpBs and Fanzors both experience the same selective pressure to maintain a high-degree of guide reprogrammability, which they overcome using similar strategies.

When comparing reRNA expression levels *in vivo* using larvae and *in vitro* using cultured cells, we found that reRNA expression was substantially higher *in vivo*. We therefore expressed the SfAV protein using the recombinant baculovirus system *in vivo* using *S. frugiperda* larvae. In addition to a his-tagged SfAV protein, we also generated a fusion construct that included an N-terminal MBP tag, as well as a C-terminal GFP-CL7 tag that allowed for visualization of protein expression. Although we could not detect the his-tag only SfAV protein, we found robust expression of the fused SfAV protein *in vivo*, with the larvae turning completely green 5 dpi (Figure E). Western-blotting using an anti-GFP antibody confirmed the expression of the full-length 145 kDa construct (Figure 5F and Supplementary Figure 11). Our results demonstrate *in vivo* expression of both the SfAV Fanzor1 and reRNA that could serve as its guides during viral infection.

### Structural analysis of Fanzors highlight key evolutionary features in the Fanzor lineage

To gain further insight into the evolution of TnpB’s and Fanzors, we performed structural analysis on AlphaFold2 (AF2) models across TnpBs and Fanzors. Overall, we observed architectural similarity between TnpB and Fanzor proteins (Figure 6A). Whereas Fanzor2s showed striking structural similarity with TnpBs, Fanzor1s were more divergent. All proteins could be visually separated into NUC and REC lobes. The REC lobe consisted of a clearly identifiable wedge domain (WED) having either a β-barrel or similar β-sheets and ɑ-helices hybrid organization, with a bundle of ɑ-helices adjacent to this structure constituting the REC1 domain. Further, the core of NUC lobe included the typical RuvC domain fold with the signature double and single ɑ-helices separated by an alternating β-strands sheet. The double ɑ-helices extended from the lobe and comprised the REC2 domain responsible for stabilization of the RNA-DNA heteroduplex in TnpB and type V CRISPR proteins (33, 34, 40, 41). On the side of the single ɑ-helix, the nucleolytic pocket with the DED catalytic triad was adjacent to the ZF domain which is likely responsible for locking the nucleic acids in the catalytic pocket (33, 40, 42). The predicted models of Fanzors also enabled the contextualization of the unusual arrangement of the catalytic site found in Fanzors and closely related TnpBs (Figure 1D). In contrast to ISDra2 TnpB structure, the predicted models reveal positioning of RuvC2 glutamate residue in a new location in the catalytic pocket that preserves positioning of the glutamate carboxylate group (Figure 6C) (40, 43).

**Figure 6.**
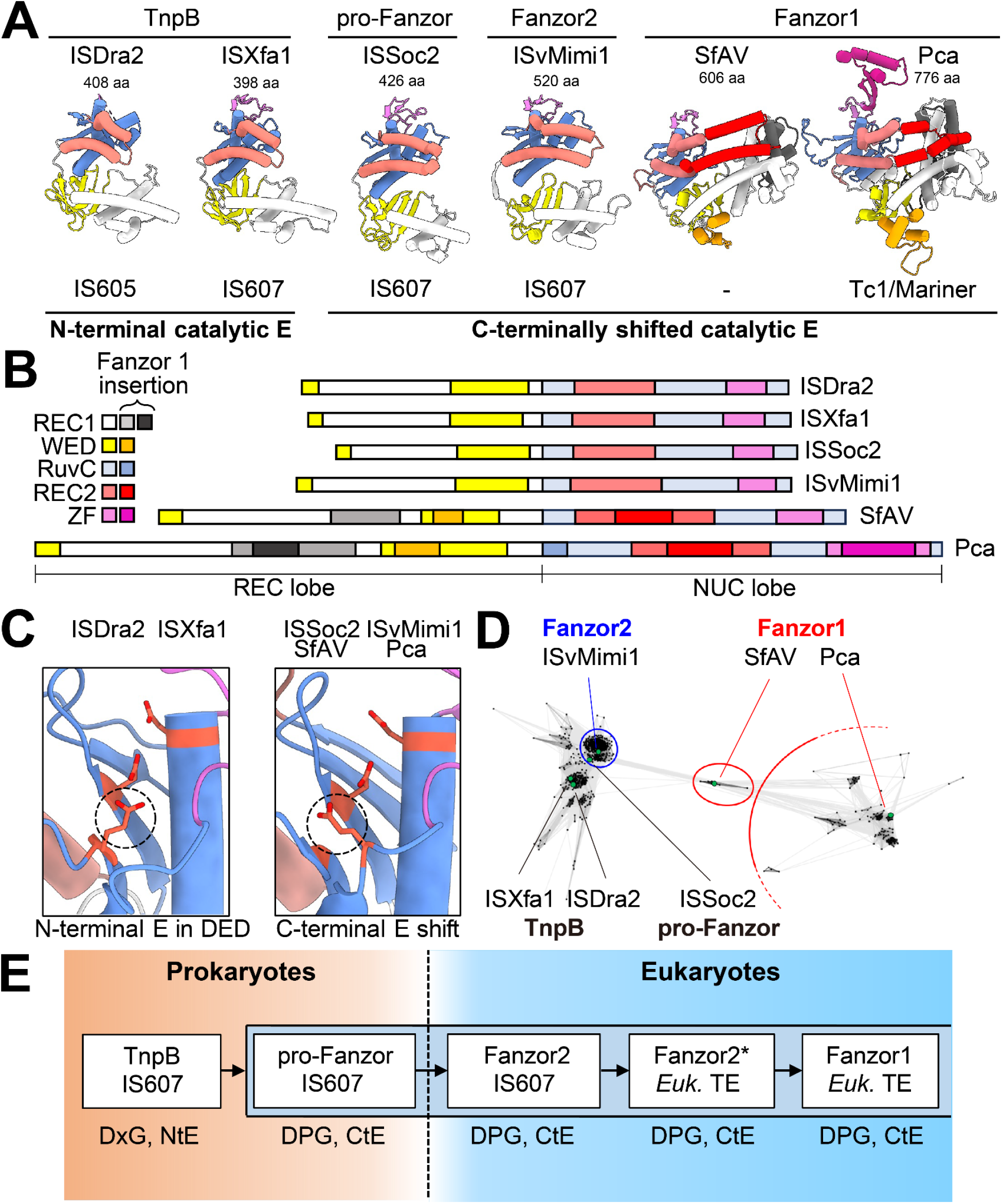
Structural comparisons of TnpBs and Fanzors highlight key evolutionary features in Fanzors. A) Example AF2 structures of TnpBs from IS200/605 and IS607 and Fanzors, highlighting different sizes of the proteins and folding of REC2 domain, which eventually binds to REC1 - a hallmark feature of Fanzor1s. B) Schematic representation of structural architecture on TnpB and Fanzor2, and regions of characteristic domain enlargement in Fanzor1. C) AF2 structures of the two types of RuvC active site organization. Dashed circle indicates the RuvC2 residue. D) CLANS clustering analysis for TnpBs and Fanzors, based on comparison of the protein sequences spanning only between RuvC1-RuvC3 catalytic residues. Note that the ascovirus/insect Fanzors are adopting an intermediate position between the cluster of Fanzor2s and other Fanzor1s. Similar arrangement is obtained with full protein sequences. E) Proposed evolutionary history from TnpB to Fanzors inferred using extant sequences.

In addition to highly conserved features, a clear trend for expansion of certain structural elements in Fanzor1s was identifiable (Figure 6A). Fanzor1s demonstrate highly similar structural features amongst themselves, suggesting a close evolutionary relationship. The hallmark structural feature of Fanzor1s is a REC2 domain comprised of a pair of kinked ɑ-helices that embraces the RNA/DNA heteroduplex, as was recently confirmed in the *Spizellomyces punctatus* Fanzor1 structure (5). Fanzor1s also feature domain insertions in the REC1 that likely interface with the heteroduplex, as well as insertions in the WED and ZF domains (Figure 6B). The expansions to the REC domains are notable, as equivalent regions in Cas12s are thought to have expanded to recognize a longer guide sequence (34). Therefore, it is conceivable that the exaggeration of these attributes in the Fanzor1 lineage, but not in prokaryotic TnpB lineages, reflects their adaptation to eukaryotes, whose larger genomes compared to prokaryotes may demand greater specificity.

Compared to Fanzor2s, the evolutionary trajectory of Fanzor1s remains less clear. In our phylogenetic analysis in Figure 2A, we found that a clade of Fanzor1s found in large DNA viruses (including SfAV) and their insect hosts represents an early diverging branch among Fanzor1s. Interestingly, pairwise similarity analysis of Fanzor sequences with CLANS independently positions this clade between Fanzor2’s and the rest of Fanzor1’s (Figure 6D). The predicted structure of SfAV suggests that these proteins bear conserved features of Fanzor1s, while lacking some insertions found in larger Fanzor1s (Figure 6A and B). Indeed, SfAV and related Fanzor1s are among the most compact Fanzor1s (550-600 residues), and their length is comparable to some Fanzor2s (∼500 residues). It is notable that among Fanzor1s, this clade of early diverging Fanzor1s bears greatest similarities to Fanzor2s, which supports a general trend of domain expansion in Fanzor1 evolution.

## DISCUSSION

Collectively, our findings suggest that an unusual type of prokaryotic TnpBs found primarily in Cyanobacteriota, the pro-Fanzors, gave rise to Fanzors in eukaryotes. Pro-Fanzors are encoded by IS607 elements, and have rare active site signatures that are conserved between pro-Fanzors and Fanzors (Figure 1C and D). As inferred through phylogenetic analysis of not only TnpBs and Fanzors, but also their associated TnpA transposases, we propose a model in which IS607 elements encoding pro-Fanzors leapt from prokaryotes to eukaryotes and their viruses, giving rise to Fanzor2s (Figure 1E). These horizontally transferred proteins were further spread and adopted by diverse TEs, as suggested by extant Fanzor2*s. We hypothesize that over long evolutionary timescales, a group of ancestral Fanzor2*s gave rise to the Fanzor1s (Figure 6E).

Our findings raise an important question as to why these pro-Fanzors were uniquely positioned to become successfully established in eukaryotes. It is unlikely that the sheer abundance of pro-Fanzors contributed to their HGT and subsequent evolution into Fanzors, as this group of TnpBs accounts for less than 1% of TnpBs in our dataset. One possible explanation could be the TnpA transposase of pro-Fanzors, as the IS607 TnpAs that mobilize dsDNA may be better suited in eukaryotes than IS200/605 TnpA that requires free ssDNA for mobility (25, 44). Whereas IS607 elements that may have recently been active can be found in eukaryotic dsDNA viruses, and sporadically in eukaryotic genomes, active eukaryotic IS200/605 elements have not been reported (7). Another possible explanation is the fact that pro-Fanzors are found in Cyanobacteriota, which have an extensive history of endosymbiosis with eukaryotes and crossdomain HGT events that include IS607 elements (6, 8, 45).

A key conserved feature across the evolution of pro-Fanzors and Fanzors is a distinct RuvC active site signature: a DPG motif at RuvC1 and a C-terminally shifted RuvC2 glutamate residue. While both of these features are independently found in additional TnpB clades, the combination of both uniquely defines pro-Fanzors and Fanzors. Recent experimental evidence suggests that the rearrangement in the RuvC2 domain decreases ssDNA *trans*-activity, a conserved biochemical feature across many Cas12s and TnpBs (4, 5, 46). How these biochemical features of these proteins contribute to their cross-domain dissemination is of great future interest and should be a subject of future investigation.

Our work illustrates the evolutionary origins of Fanzors, a widely distributed group of RNA-guided nucleases in eukaryotes. Fanzors, much like their ancestral prokaryotic TnpBs, are accessory genes of transposons. In contrast to prokaryotic TnpBs that are only found in IS200/605 and IS607 elements and their derivatives, Fanzors co-occur with a wide array of different eukaryotic transposon families, suggesting that they represent a highly successful evolutionary strategy (3, 4). Indeed, some TEs encoding Fanzors can become highly abundant in the genome, reaching up to hundreds of copies in a single genome (3). The high degree of evolutionary success of Fanzors underscores the versatility of RNA-guided endonucleases, which are ubiquitously present in all domains of life.

## DATA AVAILABILITY

All next-generation sequencing data and protein/DNA sequences generated and analyzed in this study will be made publicly available as a supplementary file.

## ACKNOWLEDGEMENTS

We thank members of the Doudna lab and the Innovative Genomics Institute for helpful discussions. We would also like to acknowledge Ms. Netravathi Krishnappa (NGS Core Operations Manager and Sequencing Specialist, Center for Translational Genomics, Innovative Genomics Institute, UC Berkeley) and Dr. Suhua Feng (The High-Throughput Sequencing Core in UCLA Broad Stem Cell Research Center) for NGS service.

## AUTHOR CONTRIBUTIONS

P.H.Y., P.S. and H.S. designed the study and performed the experiments with input and assistance with data analysis from other authors; all authors participated in manuscript and figure preparation and review.

