## Supplementary Data for "Eukaryotic RNA-guided endonucleases evolved from a unique clade of bacterial enzymes"

[1]Department of Molecular and Cell Biology, University of California, Berkeley; Berkeley, CA, USA; [2]Innovative Genomics Institute; University of California, Berkeley, CA, USA; [3]California Institute for Quantitative Biosciences (QB3), University of California, Berkeley, CA, USA; [4]Howard Hughes Medical Institute, University of California, Berkeley; Berkeley CA, USA; [5]Department of Earth and Planetary Science, University of California, Berkeley, CA, USA; [6]Department of Molecular, Cell and Developmental Biology, University of California, Los Angeles CA, USA; [7]Center for Computational Biology, University of California, Berkeley; Berkeley CA USA; [8]Howard Hughes Medical Institute, University of California, Los Angeles CA, USA; [9]Gladstone Institutes; San Francisco, CA, USA; [10]Gladstone-UCSF Institute of Genomic Immunology; San Francisco, CA, USA; [11]Molecular Biophysics and Integrated Bioimaging Division, Lawrence Berkeley National Laboratory; Berkeley, CA, USA; [12]Department of Chemistry, University of California, Berkeley; Berkeley, CA, USA;

*These authors contributed equally to the study

### SUPPLEMENTARY FIGURES

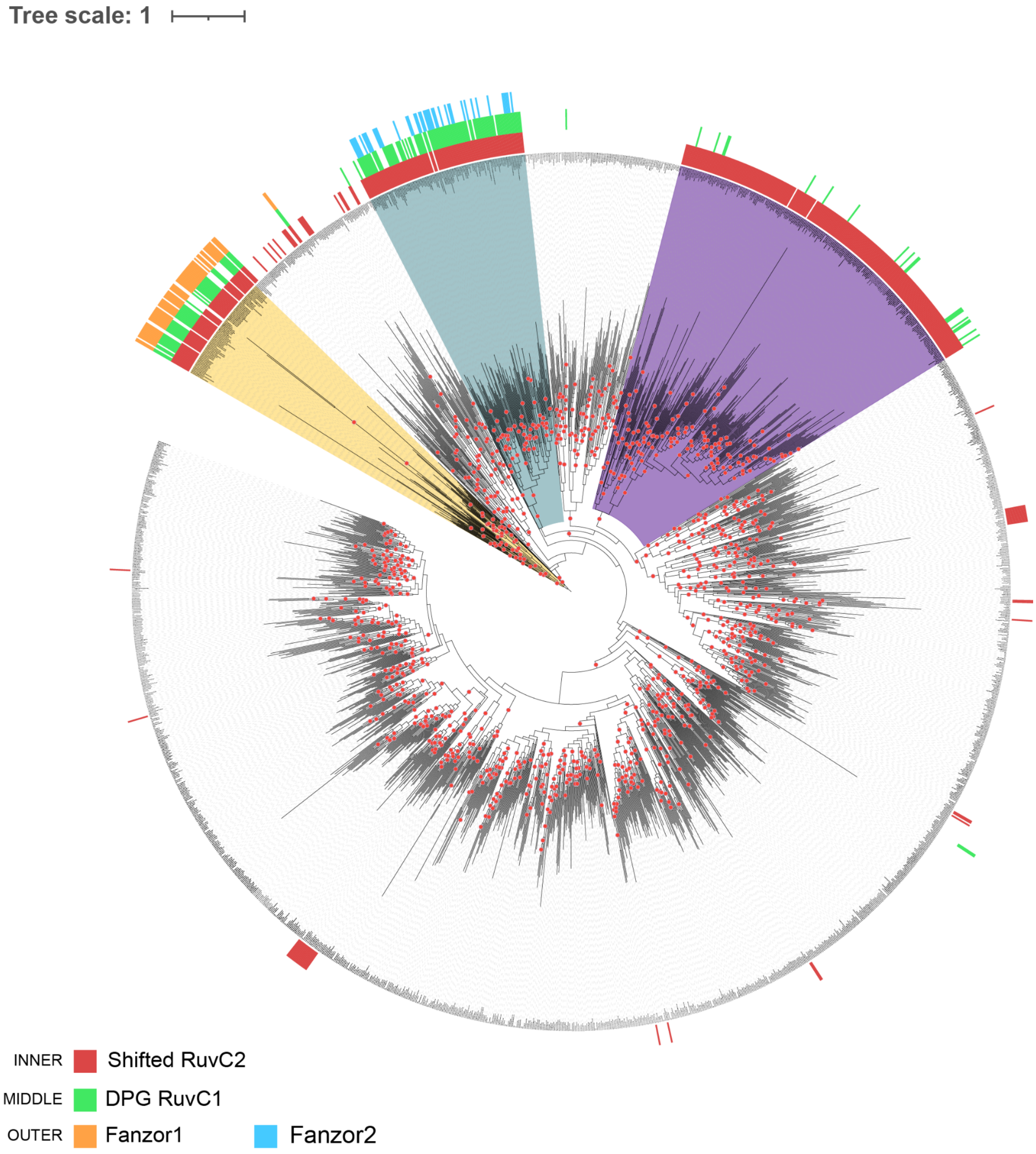


**Supplementary Figure S1. Phylogenetic tree shown of Fanzors and metagenomic TnpB set in Figure 1D.**

Red dot indicates >95 UF-bootstrap values. Yellow range shows clade containing Fanzor1, and blue range shows clade containing Fanzor2. Purple range indicates TnpBs that have independently evolved similar signature RuvC1 and RuvC2 motifs. Inner track indicates C-terminally shifted RuvC2 glutamate residue. Middle track indicates RuvC1 containing a DPG motif. Outer track annotates Fanzor1 nodes as orange, and Fanzor2 nodes as green. This tree was re-rooted using Fanzor1 containing clade as an outgroup.

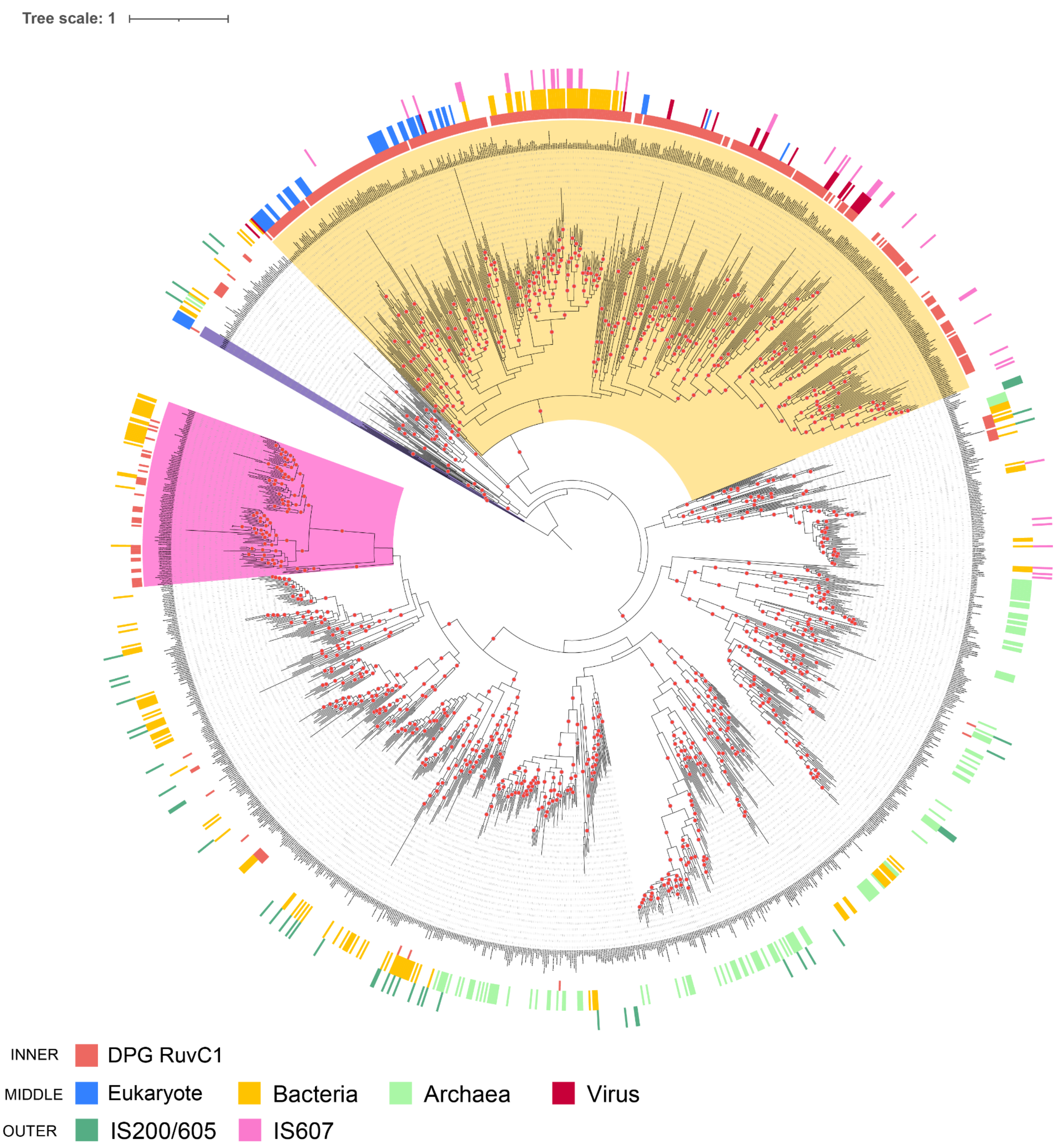


**Supplementary Figure S2. Phylogenetic tree of Fanzors and TnpBs with a shifted RuvC2 glutamate residue in Figure 1E.**

Red dot indicates >95 UF-bootstrap values. Inner track indicates the node having its RuvC1 containing a DPG motif in addition to the C-terminally shifted RuvC2 glutamate residue. Middle track annotates nodes from NCBI as being either eukaryote (blue), bacteria (yellow), archaea (light green), virus (red). Outer track annotates node as having a IS200/605 TnpA (dark green) or IS607 TnpA (pink) as its neighbor. This tree was re-rooted using Fanzor1 as an outgroup.

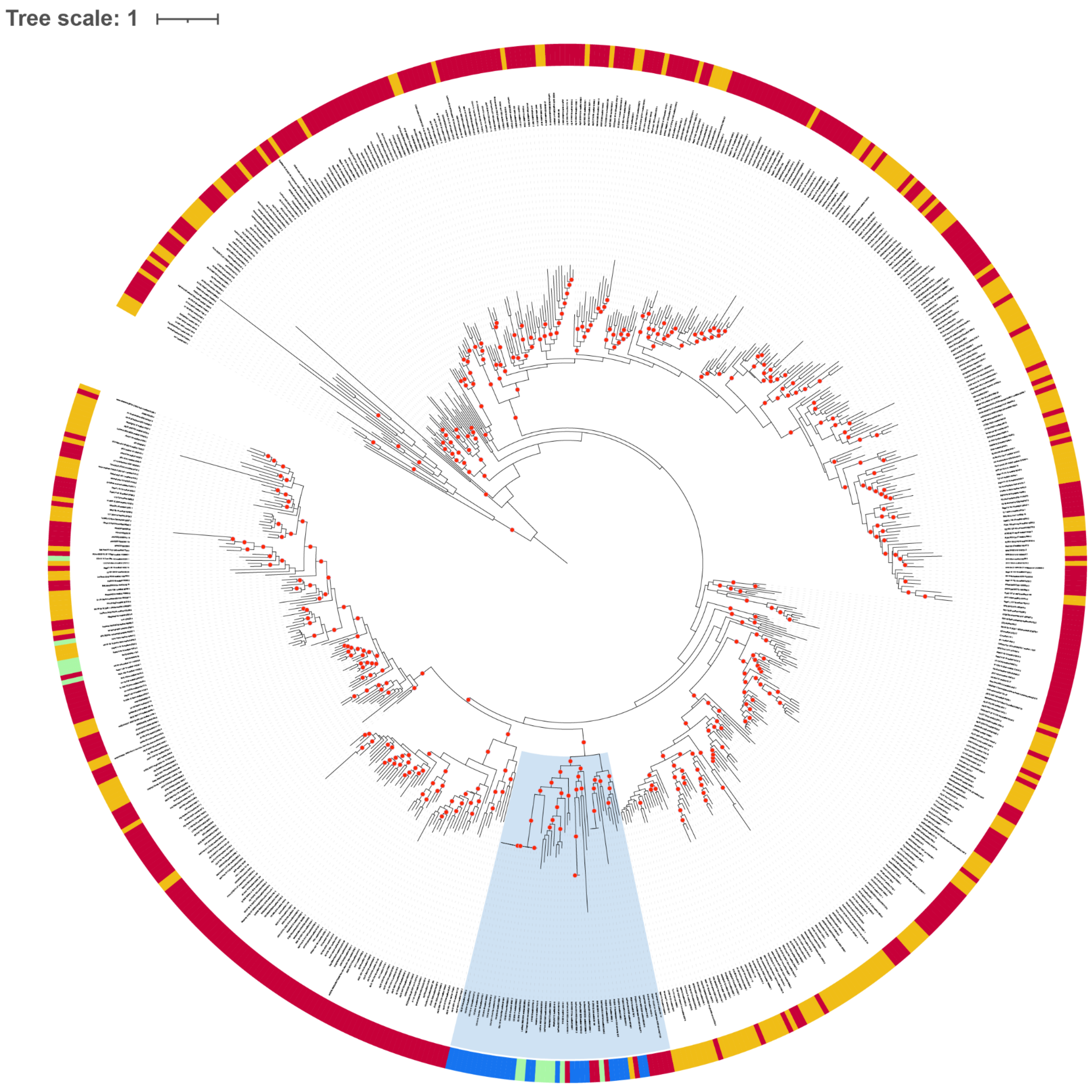


**Supplementary Figure S3. Phylogenetic tree of metagenomics derived IS607 TnpAs annotated based on their TnpB partners.**

Red dot indicates >95 UF-bootstrap values. Light blue range indicates clade containing TnpAs whose TnpB partner is a pro-Fanzor or a Fanzor. Track annotates associated TnpB as defined by RegEx shown in Supplementary Table S1 as either sharing RuvC1 and RuvC2 signatures (blue), or just RuvC2 signature (green) with Fanzors. Red indicates that the associated TnpB has a typical TnpB architecture containing the non-shifted RuvC2 residue and definitive ZF motifs as captured by the RegEx, whereas red indicates that the associated TnpB was not captured by any of the RegEx. This tree was rooted using Tn3 resolvase, which is a distant homolog of IS607 TnpA, as an outgroup.

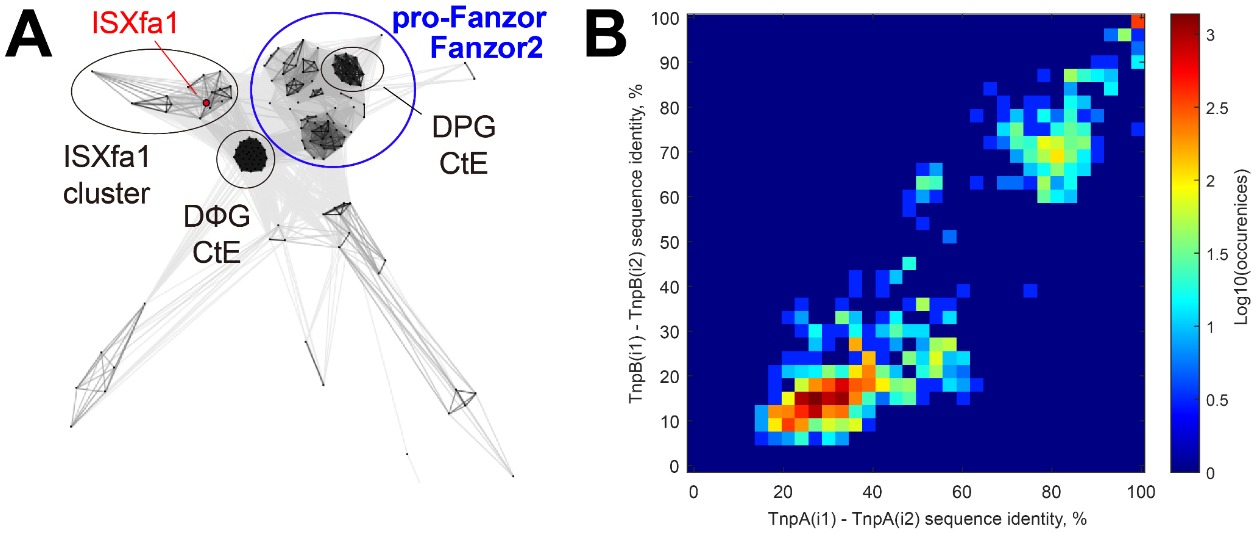


**Supplementary Figure S4. Co-conservation analysis of TnpA and TnpB/Fanzor pairs.**

A) CLANS clustering visualization of IS607-associated TnpBs from the ISFinder database, TnpBs with C-terminally shifted glutamate and DΦG motifs, and Fanzor2s. Only the TnpBs and Fanzors from the circled clusters were picked for further analysis in (B).

B) Figure shows a sequence identity histogram of pairs of TnpA and TnpB/pro-Fanzor/Fanzor2 between two distinct IS607 elements (i1 and i2) in the Fanzors-TnpB clusters. The calculated Pearson correlation coefficient 0.95 indicates high similarity of IS607 transposable elements within the selected clusters.

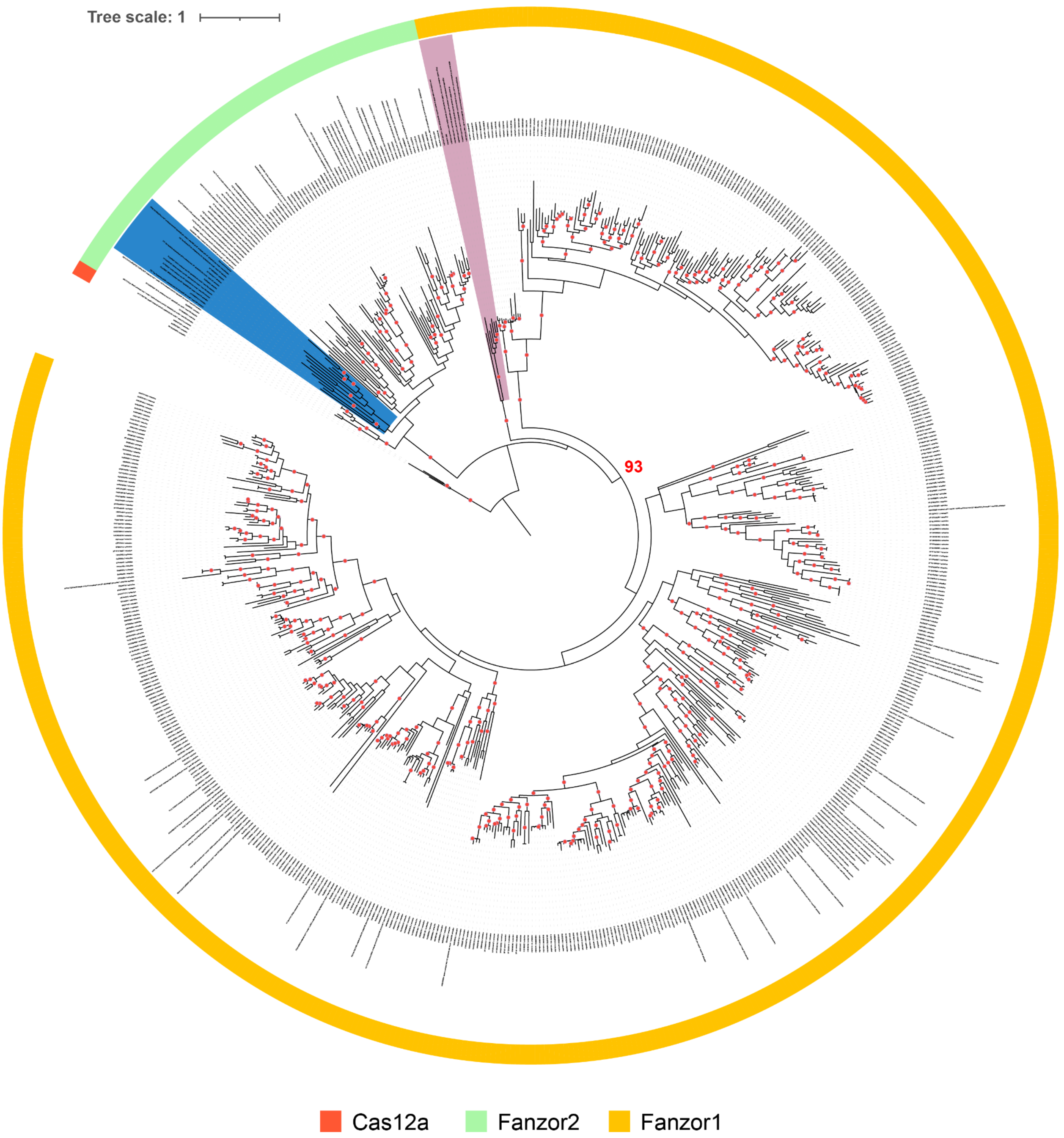


**Supplementary Figure S5. Phylogenetic tree of Fanzor sequences**

Red dot indicates >95 UF-bootstrap values. Blue range indicates clade of Fanzor2s found in molluscs and their viruses that has been co-opted by a non-IS607 transposon. Pink range indicates clade of Fanzor1s including SfAV found in lepidopterans and their viruses. Track annotates nodes as being either a Cas12a (red) Fanzor2 (green) or Fanzor1 (yellow). Tree was re-rooted using Cas12a as an outgroup. 93 indicates bootstrap values for the Fanzor1 clade.

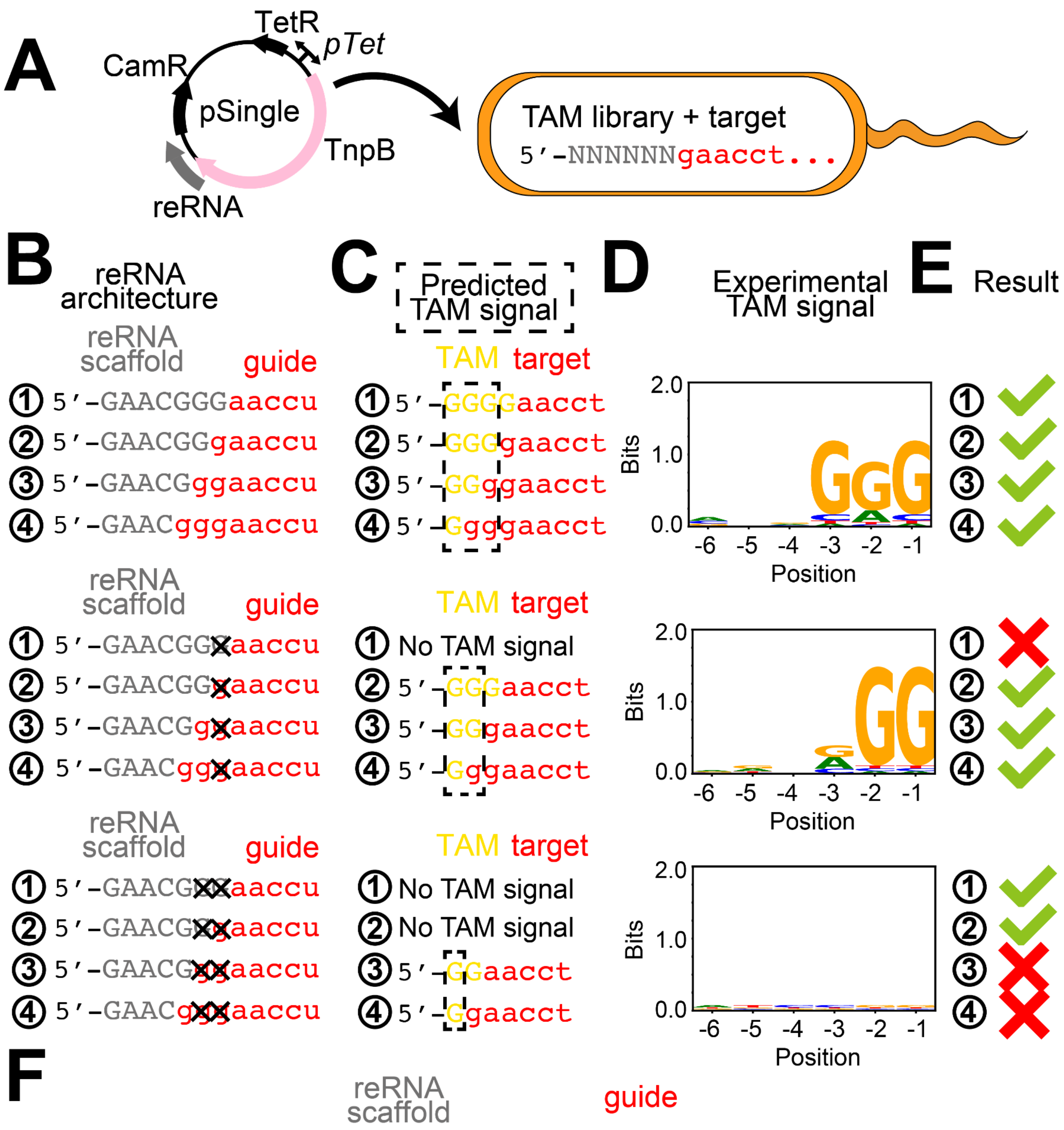


**Supplementary Figure S6. Determining the ISXfa1 reRNA boundary of the scaffold and guide.**

A) Cartoon showing the scheme of *E.coli* TAM depletion assay. The pSingle is transformed into a competent cell harboring a plasmid encoding a 6N TAM library flanking a target site. Recognition of the TAM and target site by TnpB will result in depletion of cell population bearing the TAM sequences.

B) Four hypothesized ISXfa1 reRNA boundaries of the scaffold and guide . For boundary ①, the 3' of the scaffold ends with GAACGGG, for boundary ②, the 3' of the scaffold ends with GAACGG, for boundary ③, the 3' of the scaffold ends with GAACG, and for boundary ④, the 3' of the scaffold ends with GAAC. Deleting different numbers of junctional "G" between the scaffold and guide will result in different TAM signals for different hypothesized boundaries.
C) Predicted TAM signal given different hypothesized boundaries and different deletion mutations. The TAM signals are highlighted in black box.
D) Experimental TAM motif.
E) Whether experimental results match the prediction.
F) Confirmed ISXfa1 reRNA boundary of the scaffold (gray) and guide (red).

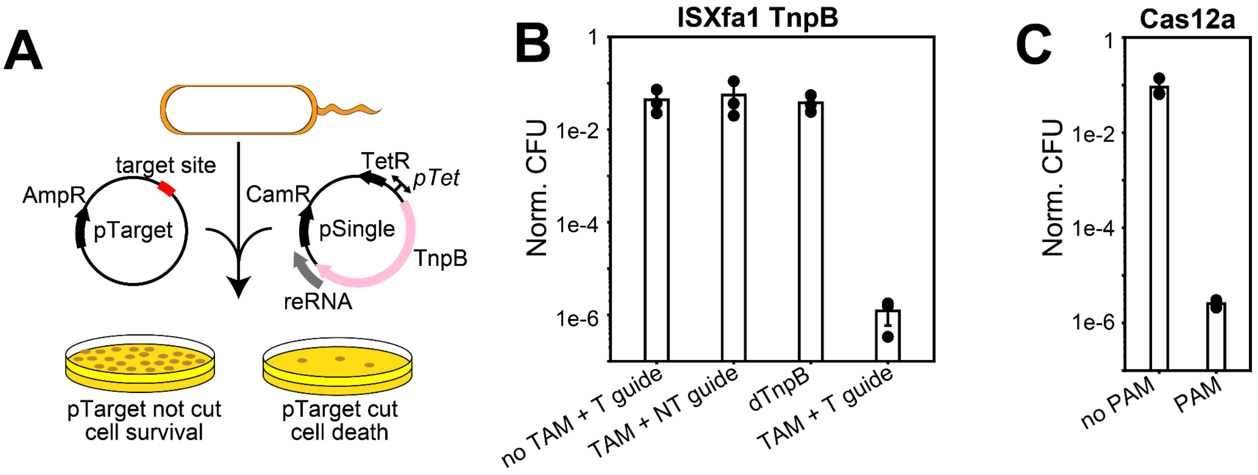


**Supplementary Figure S7. ISXfa1 TnpB plasmid interference assays in *E. coli*.**

A) Cartoon diagram depicting workflow and construct design of *E. coli* interference assay
B) Quantification of E. coli interference assay results for ISXfa1 TnpB demonstrating reRNA, TAM, and RuvC active site dependence.
C) Positive control quantifying LbCas12a interference assay results in *E. coli*, demonstrating comparable activity with ISXfa1.

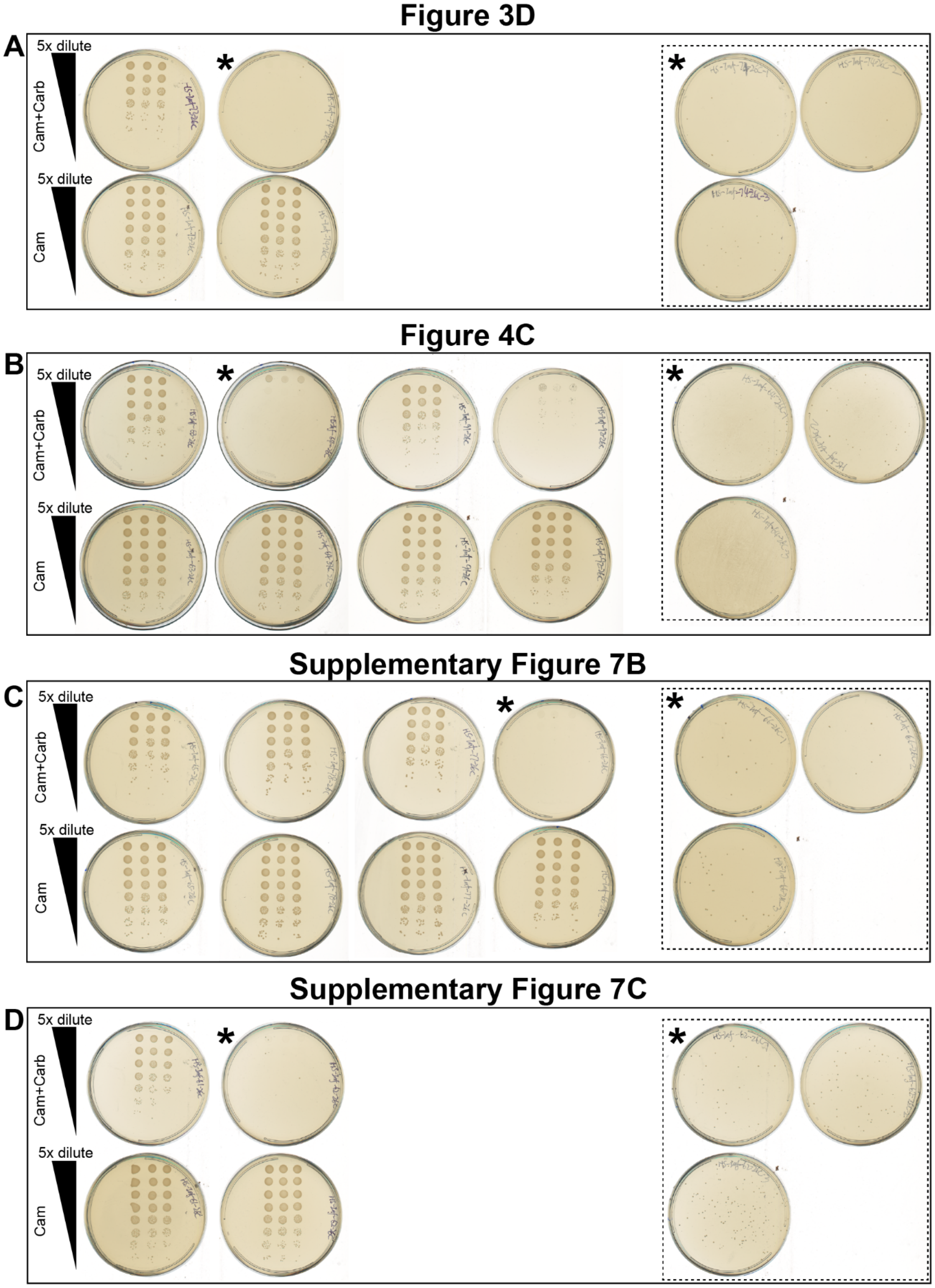


**Supplementary Figure S8. The plate images of E.coli plasmid interference assay.**
5 µL of the 5-fold serial dilution of the 1mL recovery culture after double-transformation were plated on double antibiotic LB-Agar plates (upper row) and single antibiotic plate (lower row). Each plate contains triplicate transformations. For plates with no visible colonies (asterisk), 400 µL of the original 1 mL recovery culture were plated on the double antibiotic plate (dash insets). (A) corresponds to Figure 3D, (B) to Figure 4C, (C) to Supplementary Figure S7B (D) to Supplementary S7C.

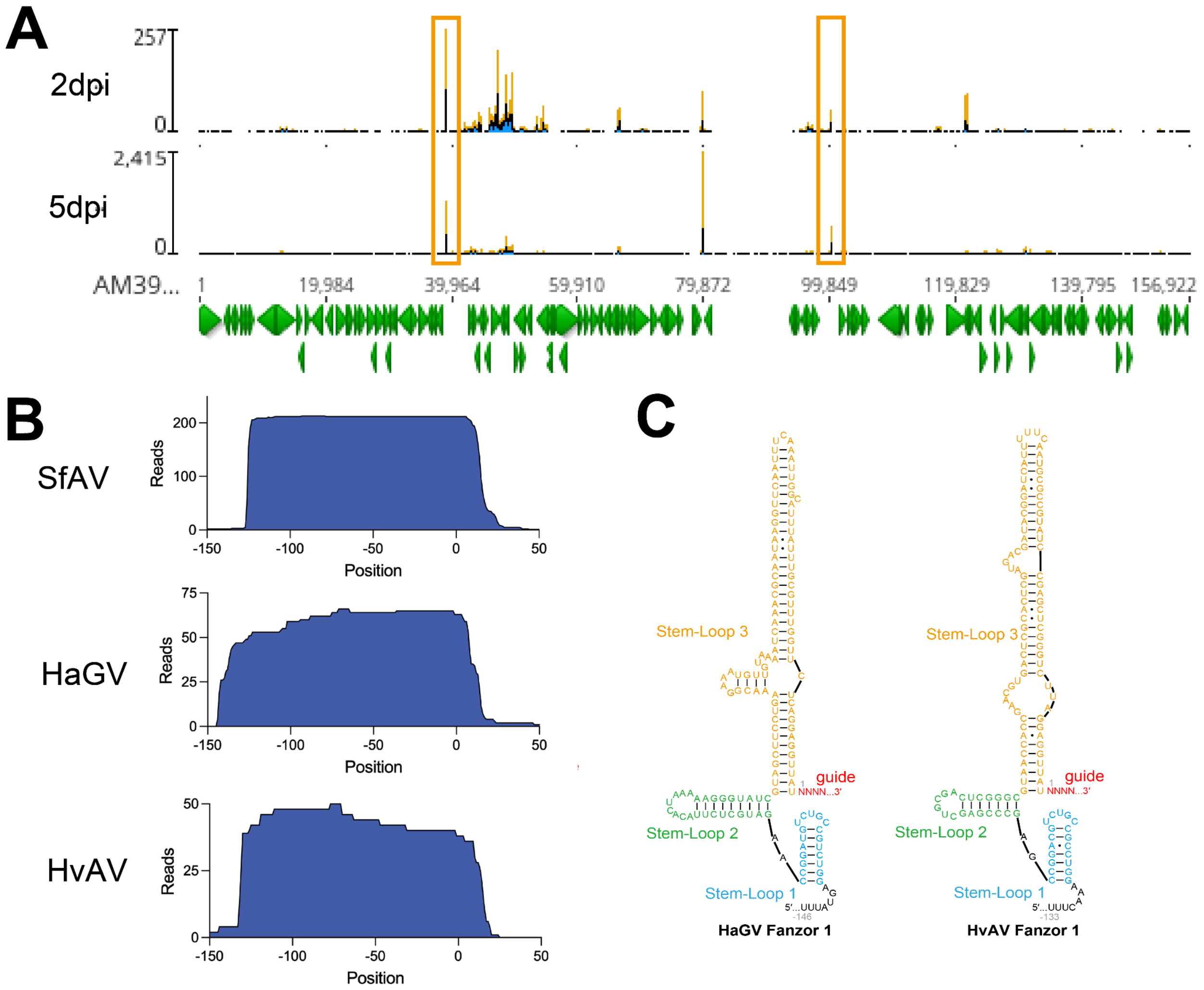


**Supplementary Figure S9. reRNA expression from insect virus Fanzors.**

A) Mapping of small RNA (<200 nt) reads from the *in vivo* expression experiment to the SfAV genome. reRNA for SfAV Fanzor is one of most highly expressed transcripts, and increased overtime from 2dpi to 5dpi.
B) Mapping of small RNA reads from Expi-Sf9 cells using recombinant baculovirus to the artificial transposon encoding locus.
C) Predicted HAgv and HVav Fanzor1 reRNA secondary structure.

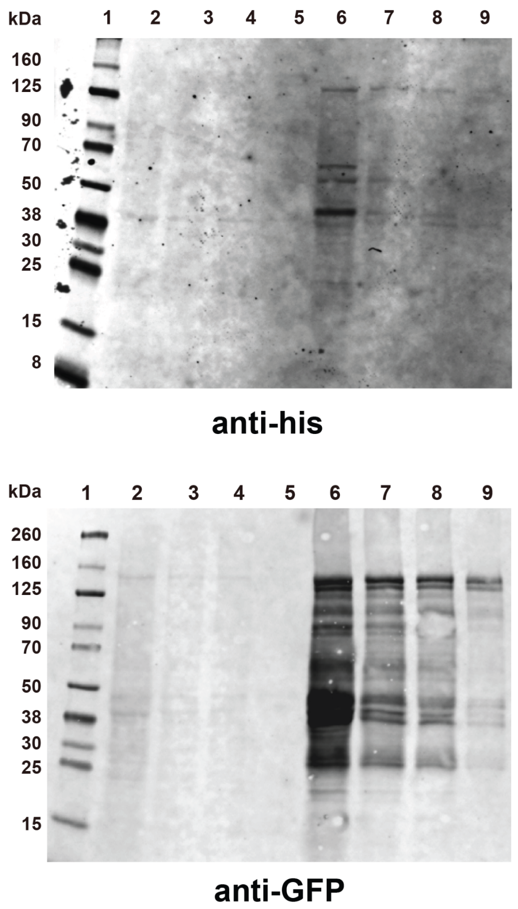


**Supplementary Figure S10. Uncropped western blot images shown in Figure 5.**
Lysate from *in vivo* expression was loaded in 5x serial dilutions. Top: anti-his primary antibody. Bottom: anti-GFP primary antibody.

### SUPPLEMENTARY TABLES

| Regular Expression | N of hits among 140k+ TnpBs |
|---|---|
| D{P}Gx(120,160)ExxxxxxCxxCx(12,18)CxxCx(4,8)D | 5053 |
| DPGx(120,160)ExxxxxxCxxCx(12,18)CxxCx(4,8)D | 1227 |
| DPGx(80,120)ExLx(40,70)CxxCx(12,18)CxxCx(4,8)D | 171 |
| D{P}Gx(80,120)ExLx(40,70)CxxCx(12,18)CxxCx(4,8)D | 62063 |

**Supplementary Table S1. Regular Expression used to categorize TnpB sequences into different categories.**

| Accession | Reported Species | Phlya |
|---|---|---|
| BAY47071.1 | Scytonema sp. HK-05 | Cyanobacteriota |
| OQX24300.1 | Desulfobacteraceae bacterium IS3 | Desfulobacteria |
| WP 242056789.1 | unclassified Planktothrix | Cyanobacteriota |
| WP 236117061.1 | Hassalia byssoidea | Cyanobacteriota |
| TAD80975.1 | Oscillatoriales cyanobacterium | Cyanobacteriota |
| WP 015227666.1 | Halothece sp. PCC 7418 | Cyanobacteriota |
| WP 235622542.1 | Nostocaceae | Cyanobacteriota |
| BAB78230.1 | Nostoc sp. PCC 7120 = FACHB-418 | Cyanobacteriota |
| WP 041780216.1 | Allocoleopsis franciscana | Cyanobacteriota |
| WP 071108120.1 | Moorena producens | Cyanobacteriota |
| WP 229420144.1 | Moorena | Cyanobacteriota |
| WP 245587445.1 | Cylindrospermum stagnale | Cyanobacteriota |
| TAE02002.1 | Oscillatoriales cyanobacterium | Cyanobacteriota |
| WP 127081850.1 | Dulcicalothrix desertica | Cyanobacteriota |
| WP 233154214.1 | Scytonema sp. HK-05 | Cyanobacteriota |
| WP 190628969.1 | unclassified Calothrix | Cyanobacteriota |
| WP 263747204.1 | Plectonema radiosum | Cyanobacteriota |
| WP 168567848.1 | Oxynema aestuarii | Cyanobacteriota |
| WP 096692845.1 | unclassified Calothrix | Cyanobacteriota |
| WP 272819120.1 | Scytonema hofmannii | Cyanobacteriota |
| KAB8314710.1 | Tolypothrix campylonemoides VB511288 | Cyanobacteriota |
| WP 137985626.1 | Moorena producens | Cyanobacteriota |
| WP 008177940.1 | Moorena | Cyanobacteriota |
| WP 263012505.1 | unclassified Laspinema | Cyanobacteriota |
| TAD89774.1 | Oscillatoriales cyanobacterium | Cyanobacteriota |
| TAE00453.1 | Oscillatoriales cyanobacterium | Cyanobacteriota |
| WP 265233693.1 | Lyngbya sp. CCAP 1446/10 | Cyanobacteriota |
| WP 248277801.1 | Brasilonema sp. UFV-L1 | Cyanobacteriota |
| QDL12051.1 | Brasilonema sennae CENA114 | Cyanobacteriota |
| TAE61799.1 | Nostocales cyanobacterium | Cyanobacteriota |
| WP 197285365.1 | Planktothricoides sp. SR001 | Cyanobacteriota |
| EKD09239.1 | Arthrospira platensis C1 | Cyanobacteriota |

| | | |
|---|---|---|
| WP 099887212.1 | Synechococcus sp. 63AY4M2 | Cyanobacteriota |
| PIK98325.1 | Synechococcus sp. 63AY4M1 | Cyanobacteriota |
| WP 265415802.1 | Aphanothece sacrum | Cyanobacteriota |
| WP 242060420.1 | Planktothrix sp. FACHB-1375 | Cyanobacteriota |
| WP 105220190.1 | Gloeocapsopsis dulcis | Cyanobacteriota |
| WP 236117273.1 | Hassalia byssoidea | Cyanobacteriota |
| WP 242060286.1 | Planktothrix sp. FACHB-1375 | Cyanobacteriota |
| AFY80297.1 | Oscillatoria acuminata PCC 6304 | Cyanobacteriota |
| WP 231296431.1 | Arthrospira platensis | Cyanobacteriota |
| EKD07988.1 | Arthrospira platensis C1 | Cyanobacteriota |
| AFZ27794.1 | Cylindrospermum stagnale PCC 7417 | Cyanobacteriota |
| WP 199341614.1 | Nostocales | Cyanobacteriota |

**Supplementary Table S2. Prokaryotic relatives of Fanzors and their NCBI accession ID and reported species showing enrichment of Cyanobacteriota in this clade.**
